# Inhibition of IRAK4 by microbial trimethylamine blunts metabolic inflammation and ameliorates glycemic control

**DOI:** 10.1101/277434

**Authors:** Julien Chilloux, Francois Brial, Amandine Everard, David Smyth, Petros Andrikopoulos, Liyong Zhang, Hubert Plovier, Antonis Myridakis, Lesley Hoyles, José Maria Moreno-Navarrete, Jèssica Latorre Luque, Viviana Casagrande, Rosella Menghini, Blerina Ahmetaj-Shala, Christine Blancher, Laura Martinez-Gili, Selin Gencer, Jane F. Fearnside, Richard H. Barton, Ana Luisa Neves, Alice R. Rothwell, Christelle Gérard, Sophie Calderari, Mark J. Williamson, Julian E. Fuchs, Lata Govada, Claire L. Boulangé, Saroor Patel, James Scott, Mark Thursz, Naomi Chayen, Robert C. Glen, Nigel J. Gooderham, Jeremy K. Nicholson, Massimo Federici, José-Manuel Fernández-Real, Dominique Gauguier, Peter P. Liu, Patrice D. Cani, Marc-Emmanuel Dumas

## Abstract

The global type 2 diabetes epidemic is a major health crisis and there is a critical need for innovative strategies to fight it. Although the microbiome plays important roles in the onset of insulin resistance (IR) and low-grade inflammation, the microbial compounds regulating these phenomena remain to be discovered. Here, we reveal that the microbiome inhibits a central kinase, eliciting immune and metabolic benefits. Through a series of *in vivo* experiments based on choline supplementation, blocking trimethylamine (TMA) production then administering TMA, we demonstrate that TMA decouples inflammation and IR from obesity in the context of high-fat diet (HFD) feeding. Through *in vitro* kinome screens, we reveal TMA specifically inhibits Interleukin-1 Receptor-associated Kinase 4 (IRAK4), a central kinase integrating signals from various toll-like receptors and cytokine receptors. TMA blunts TLR4 signalling in primary human hepatocytes and peripheral blood monocytic cells, and improves mouse survival after a lipopolysaccharide-induced septic shock. Consistent with this, genetic deletion and chemical inhibition of IRAK4 result in similar metabolic and immune improvements in HFD. In summary, TMA appears to be a key microbial compound inhibiting IRAK4 and mediating metabolic and immune effects with benefits upon HFD. Thereby we highlight the critical contribution of the microbial signalling metabolome in homeostatic regulation of host disease and the emerging role of the kinome in microbial–mammalian chemical crosstalk.

## Main

Globally, diabetes affects 529 million individuals^1^, claiming 1.6 million lives per year according to the World Health Organization. Insulin resistance (IR) is a multifactorial condition and now increasingly common due, among other factors, to the increase in prevalence of a sedentary lifestyle, Western-style foods and obesity. IR is a risk factor for developing type 2 diabetes and cardiovascular diseases^2^. One of the hallmarks of these disorders is the early development of a chronic, systemic low-grade inflammation^3^. The interaction between high-fat diet (HFD) feeding and the gut microbiome has a strong impact on the onset of IR^4–6^: bacterial lipopolysaccharides (LPS) and dietary fats trigger low-grade inflammation^2^ through activation of *Toll*-like receptor 4 (TLR4), a process called metabolic endotoxemia^7^.

This is supported by a complex communication that occurs between the gut microbiota and the innate immune system^8^, with consequences on metabolic homeostasis^9^. Whilst some of the functional signalling molecules mediating microbial–mammalian chemical crosstalk have been characterized, a limited number of metabolite classes and their targets have been identified (G protein-coupled^10^ and nuclear receptors^11^). It is, however, hypothesized that microbial metabolites potentially affect other pharmacological target classes such as kinases^12^. In previous studies, we and others have identified a family of microbiome-associated metabolites, methylamines, associated with IR, non-alcoholic fatty liver disease^13^ and atherosclerosis^14^, but their mechanisms of action on mammalian hosts remain unclear. Trimethylamine (TMA) is one of the most abundant microbial metabolites produced by the gut microbiota. We first reported that TMA may be associated with IR^12^. TMA results from microbial metabolism of dietary choline, carnitine and trimethylamine *N*-oxide (TMAO)^15–19^ before being absorbed and *N*-oxidized into TMAO by hepatic flavin-containing mono-oxygenase 3 (FMO3)^20^. After initial reports associating TMAO with adverse cardiovascular outcomes^13,16^, it has since emerged that FMO3 inactivation was beneficial for several metabolic outcomes^21–24^, strongly suggesting that TMA and TMAO have distinct biological roles.

Here, we decipher the role of TMA in the microbiota–host kinome chemical crosstalk in IR through i) identification of gut-derived microbial metabolites associated with HFD-induced impaired glucose tolerance (IGT), IR and obesity, ii) pharmacological target screening of discriminant microbial metabolites, and iii) mechanistic validation of the pathophysiological relevance of pharmacological interactions with *in vitro* and *in vivo* models. Using this approach, we discovered a novel mechanism by which gut microbial TMA acts as an IRAK4 inhibitor and directly improves the host immune and metabolic tone.

### Lack of hepatic inflammation in HFD and the choline supplementation hypothesis

We initially carried out a longitudinal pathophysiological monitoring in a cohort of C57BL/6 mice (n=5 for HFD and n=6 for chow diet (CHD)) rapidly developing obesity and IGT when fed a 65% kCal HFD with a range of macronutrient and micronutrient differences compared to CHD (**Extended Data Fig. 1a-b, Fig. 1a** and **Supplementary Table 1**). Liver transcriptomics (n=5-6 per diet group) identified 2,084 significantly (FDR < 0.1) differentially expressed genes between CHD-fed and HFD-fed mice (**Extended Data Fig. 1c, Fig. 1b** and **Supplementary Table 1**). Gene ontology and signalling pathway impact analyses (**Extended Data Fig. 2, Supplementary Table 2-4**) demonstrated upregulation of protein processing in the endoplasmic reticulum, whereas carbohydrate metabolism, circadian rhythm and AMPK signalling were significantly inhibited, consistent with existing literature^25,26^. Surprisingly, inflammation-associated pathways were not differentially affected: expression of genes coding for acute phase serum amyloid A proteins *Saa1* and *Saa2* upregulated in response to acute inflammation and associated with type 2 diabetes was reduced, whereas expression of pro-inflammatory cytokines such as *Il6* and *Il1β* was not significant (**Fig. 1b** and **Supplementary Table 4**). This is in contrast to established knowledge as low-grade inflammation is one of the key features associated with IR^2,6^ and HFD^27^. Beyond % of fat content as a macronutrient, we therefore looked for variation in micronutrients and microbial metabolites that may contribute to this phenotype by performing metabolic profiling of the urinary metabolome of this mouse cohort (CHD and HFD, 4 time-points) by ^1^H-NMR spectroscopy followed by multivariate modelling. Our analysis showed diet is the main factor influencing the metabolic profiles, followed by age (**Extended Data Fig. 1d**); the model’s goodness-of-prediction parameters being highly significant upon permutation testing (**Extended Data Fig. 1e**). The metabolic signature of HFD-feeding compared to CHD was mainly associated with TMA excretion (**Fig. 1c**), consistent with our previous reports using this diet^13,28^. We used variance components analysis to decompose the contribution of diet and age to the excretion of these three metabolites, showing that the HFD contribution overwhelms the contribution of age (**Fig. 1d**). Since the HFD contains >15 g of choline per kg of diet, we then investigated whether supplementation in this micronutrient could modulate the metabolic and immune variation in response to HFD feeding.

**Figure 1.**
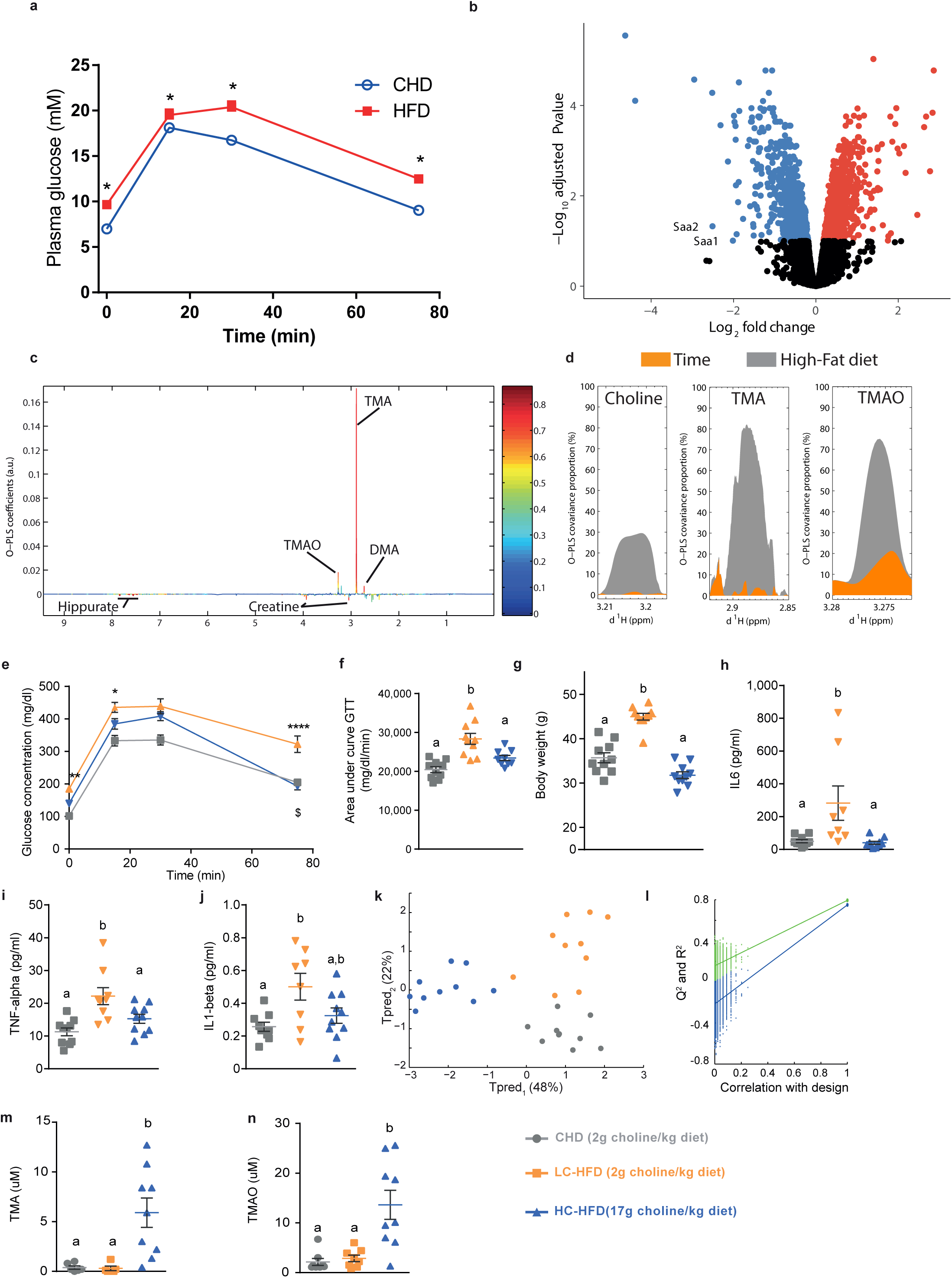
Choline supplementation improves glucose homeostasis and inflammation after 5 months of HFD. Male mice were weaned at 3 weeks and fed a CHD diet, before being randomly assigned to CHD group (n=68) or HFD (n=47) groups and monitored at 5 months of age. Effect of HFD-feeding on (**a**) glucose tolerance test (GTT). (**b**) Volcano plot of differentially expressed liver genes, highlighting that *Saa1* and *Saa2* are down-regulated in HFD after 4 months of HFD. Red dots, significantly (FDR < 0.1) upregulated genes; blue dots, significantly (FDR < 0.1) downregulated genes. (**c**) Metabolic signature of diet on O-PLS model coefficient plot; line plot corresponds to covariance and the color-scale represents proportion of variance explained. (**d**) Assessment of the contribution of factors diet and age on choline, TMA and TMAO by O-PLS variance components analysis. 5-week-old C57BL/6J mice were fed a CHD diet, a low-choline HFD or a low-choline HFD supplemented with choline (n=10) and were phenotyped after 5 months. (**e**) Plasma glucose concentration during an ipGTT. (**f**) Area under the curve of the plasma glucose concentration during an ipGTT. (**g**) Body weight (BW) measured after 5 months of diet. (**h-j**) Quantification of circulating cytokines: IL6 (**h**), TNFα (**i**) and IL1β (**j**). (**k**) Scores plot from a cross-validated O-PLS-DA model segregating the three groups of mice according to diet based on isotopic quantification of methylamines by UPLC-MS/MS, each component’s explained variance is shown in parenthesis. (**l**) Empirical assessment of the significance of O-PLS goodness-of-fit parameters by generating a null distribution with 10,000 random permutations. (**m-n**) Quantification of circulating TMA (**m**) and TMAO (**n**) by UPLC-MS/MS. Data are means ± s.e.m. One-way ANOVA followed by Tukey’s *post hoc* tests (superscript letters for factor levels *P*<0.05) on log-transformed data except (**k&l**).

### Choline supplementation corrects HFD-induced inflammation and IR

To test the effect of choline supplementation itself on glucose tolerance and IR in an HFD context, we initiated a series of *in vivo* studies. We first fed C57BL/6 mice with a 65% kCal HFD containing 2 g/kg or 17 g/kg of choline (i.e.: LC-HFD and HC-HFD, respectively; **Fig. 1e-n**). To evaluate metabolic homeostasis, we performed glucose tolerance tests (GTTs), which displayed a similar pattern with normalisation of cumulative glycemia on HC-HFD compared to LC-HFD (**Fig. 1e-f**). Body weight (BW) was significantly increased in mice fed a LC-HFD from weaning until 5 months of age compared to mice fed a chow diet and remarkably compared to mice fed HC-HFD (**Fig. 1g**), which was not related to food intake (**Extended Data Fig. 3a**). We then profiled circulating pro-inflammatory cytokines such as IL6, IL1*β*, TNFα (**Fig. 1h-j**) and phosphorylation of NF-κB regulating their transcription (**Extended Data Fig. 3b,c**), which was also suggestive of an improvement of HFD-induced inflammation by choline supplementation.

### Targeted analysis of choline and methylamine pathways

To document the effect of choline supplementation on choline-related metabolic pathways, we further refined our isotopic UPLC-MS/MS quantification^29,30^ to evaluate plasma concentrations of choline- and carnitine-derived metabolites leading to TMA and TMAO (**Extended Data Fig. 3d,e**). An O-PLS-DA model significantly segregates the three treatment groups at 5 months of age (i.e., 4 months of HFD) and highlights a choline supplementation effect on the first predictive component (**Fig. 1k,l**). Quantifications all reflect an increase in TMA and TMAO in the HC-HFD in line with choline supplementation (**Fig. 1m,n**). In particular, the circulating TMA levels were similar in the CHD and the LC-HFD (0.38 µM in CHD vs. 0.3 µM in LC-HFD) and were about 20 times lower than in the HC-HFD (5.9 µM).

These results depict an increased microbial conversion of choline into TMA in our HC-HFD, an observation already made in a previous study^13^. These results suggest that TMA could mediate the metabolic and immune benefits of choline supplementation.

### Baseline metabolic phenotyping after 2-month choline modulation in HFD

To further assess the impact of choline supplementation on metabolic homeostasis and low-grade inflammation in HFD contexts, we performed a second experiment, feeding C57BL/6 mice a 60% kcal HFD with a time-frame comparable to the ones used in subsequent experimental designs using osmotic minipumps for constant subcutaneous TMA delivery. GTTs and weekly BWs showed a similar trend after 8 weeks of HFD-feeding (**Fig. 2a-d**). Choline supplementation not only improved glucose tolerance (**Fig. 2a,b**) but also the Matsuda index (**Fig. 2c**), a marker for IR derived from oral GTTs^31^. The normalisation of insulin sensitivity was confirmed through insulin tolerance tests (ITTs) and insulin-induced hepatic Akt phosphorylation assays (**Fig. 2e-h**). Notably, the normalisation of insulin-induced hepatic AKT phosphorylation assays was primarily associated with a reduction of basal level of P-AKT in HC HFD compared to LC HFD observed in saline condition demonstrating the reduction of IR (**Fig. 2g**). Indeed, it has been previously demonstrated that HFD increased the basal levels of P-AKT and this participates to IR^32^. We further characterized that HFD feeding increased hepatic NF-κB phosphorylation compared to CHD, which was normalised by choline supplementation (**Fig. 2i,j**), a pattern that was also observed for expression of acute-phase proteins that are typically upregulated in response to tissue inflammation^33^ – *Saa1*, *Saa2* and *Saa3* (**Fig. 2k-m**).

**Figure 2.**
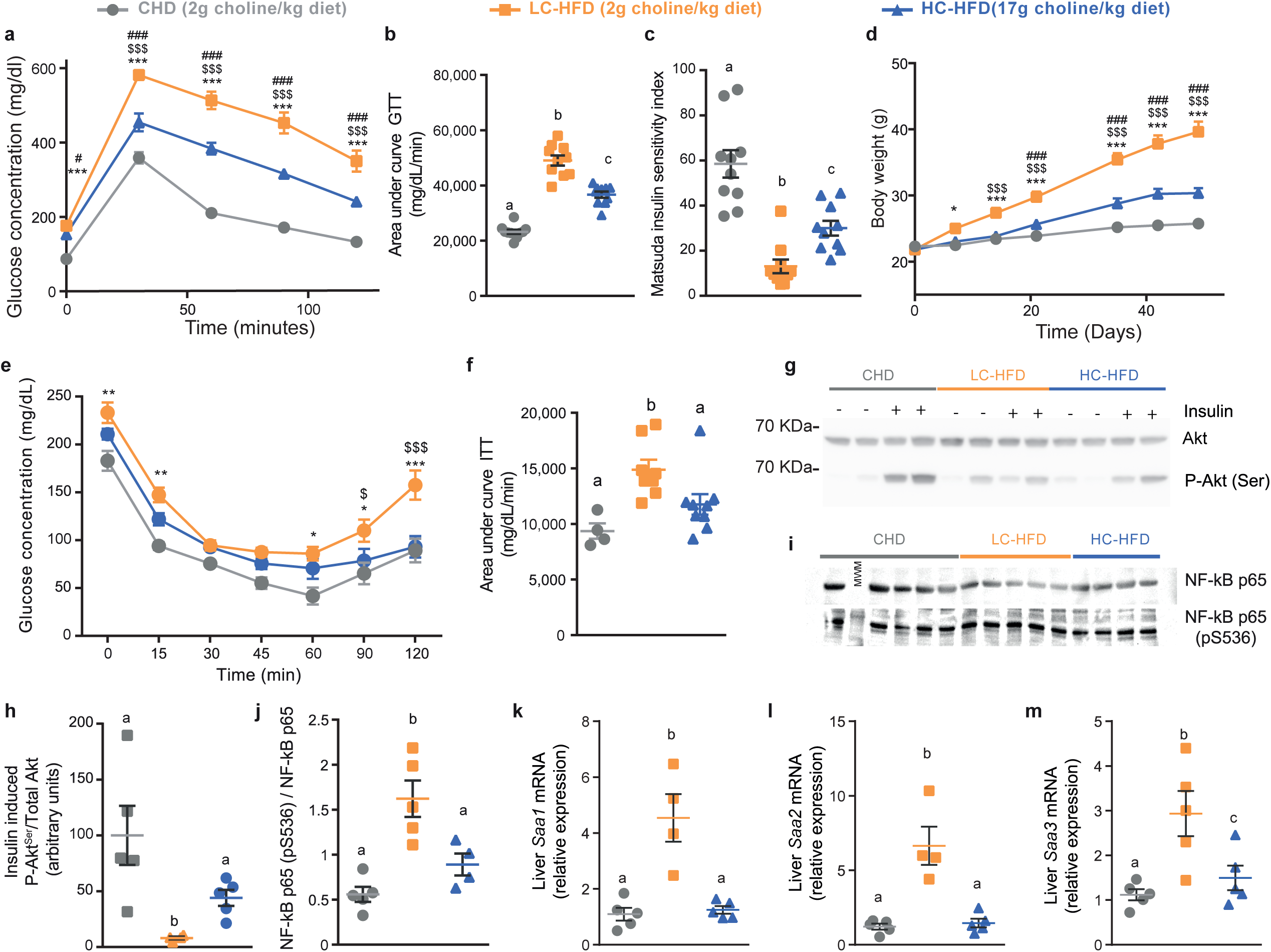
Choline supplementation corrects HFD adverse effects on glucose homeostasis, insulin sensitivity and inflammation after 8 weeks of diet. Five-week-old C57BL/6J mice were fed a CHD diet, a low-choline HFD or a low-choline HFD supplemented with choline (n = 10) and were phenotyped after 8 weeks. (**a**) Plasma glucose concentration during an ipGTT. (**b**) Area under the curve of the plasma glucose concentration during an ipGTT. (**c**) Matsuda insulin sensitivity index calculated from the ipGTT. (**d**) Body weight measured periodically during 8 weeks of diet. (**e**) Plasma glucose concentration during an ipITT. (**f**) Area under the curve of the plasma glucose concentration during an ipITT. (**g-j**) Western-blot analysis of liver Akt phosphorylation (**g-h**) and NF-kB phosphorylation (**i-j**). (**k-m**) Expression of hepatic acute phase inflammation proteins markers *Saa1* (**k**), *Saa2* (**l**) and *Saa3* (**m**). Data are means ± s.e.m. (**c**, **d**, **f**, **h-m**) One-way ANOVA followed by Tukey’s *post hoc* tests (superscript letters for factor levels *P*<0.05) on log-transformed data. (**a**, **b**, **e**) Non-parametric two-sided Mann-Whitney test was used for each timepoint, significance comparison signs are as follow: *=CHD vs LC-HFD, $=LC-HFD vs HC-HFD and #=CHD vs HC-HFD.

### Chemical blockage of bacterial TMA biosynthesis and loss of metabolic benefits

To test whether the beneficial effects of choline supplementation are mediated by its microbial product TMA, we sought to block bacterial TMA production in HC-HFD-fed mice both non-specifically and specifically using a wide-spectrum antibiotics cocktail or 3,3-dimethyl-1-butanol (DMB), inhibiting microbial TMA-lyase^34^, respectively. We first confirmed the functional blockage of TMA production in mice fed a HC-HFD, resulting in a drastic drop in circulating and excreted TMA and TMAO following 1% DMB administration (**Extended Data Fig. 4a-d**). In accordance with our hypothesis, both antibiotic and DMB treatments abolished the effects of choline supplementation-induced improvements in HFD, in particular for glucose tolerance, insulin sensitivity (as suggested by the Matsuda index, **Fig. 3a-c**, **Extended Data Fig. 4e,f**) and hepatic insulin sensitivity as shown by the absence of a beneficial effect of choline supplementation on insulin-induced Akt phosphorylation (**Fig. 3d-e**). The metabolic improvements associated with chemical TMA synthesis blockage were not due to changes in BW gain (**Extended Data Fig. 4e**), thereby strongly suggesting that the effects on glucose metabolism induced by inhibiting bacterial TMA production in HC-HFD were decoupled from obesity.

**Figure 3.**
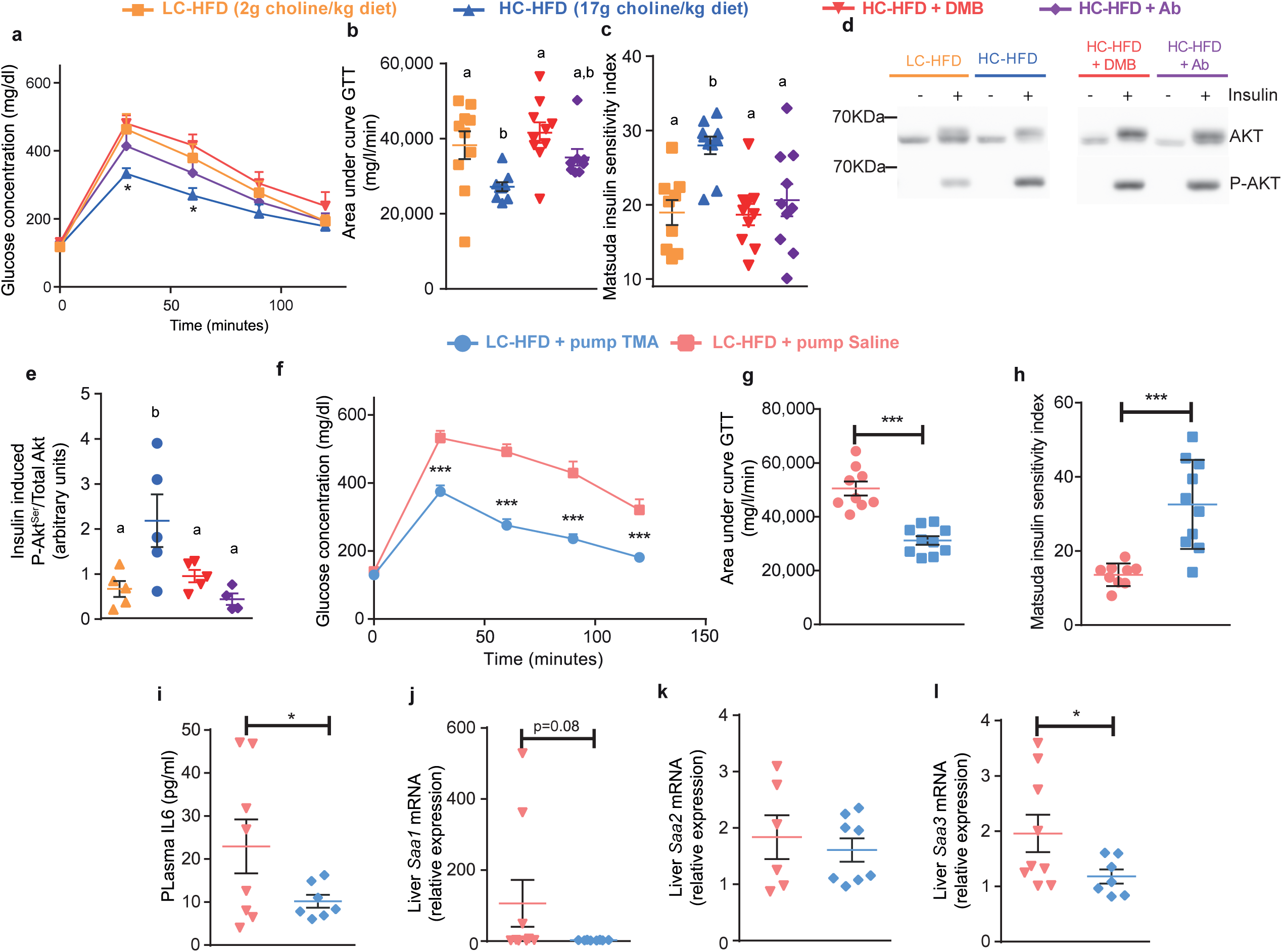
Blocking TMA production from choline cancels metabolic benefits from choline supplementation and chronic TMA chronic treatment mimics it. (**a-e**) Five-week-old C57BL/6J mice were fed a low-choline HFD or a low-choline HFD supplemented with choline (n = 10). Two strategies were used in parallel to block TMA production from choline by the gut microbiota using either 1% DMB in the diet or an antibiotic treatment. (**a**) Plasma glucose concentration during an ipGTT. (**b**) Area under the curve of the plasma glucose concentration during an ipGTT. (**c**) Matsuda insulin sensitivity index calculated from the ipGTT. (**d-e**) Western-blot analysis of liver Akt phosphorylation. (**f-l**) Mice were weaned at 3 weeks and fed a low-choline HFD before being fitted with osmotic minipumps delivering a chronic circulating dose of TMA (0.1 mM) for 6 weeks (n = 9 saline HFD, n = 8 TMA HFD). (**f**) Plasma glucose concentration during an ipGTT. (**g**) Area under the curve of the plasma glucose concentration during an ipGTT. (**h**) Matsuda insulin sensitivity index calculated from the ipGTT. (**i**) Quantification of circulating IL6. (**j-l**) Expression of hepatic acute-phase inflammation protein markers *Saa1* (**j**), *Saa2* (**k**) and *Saa3* (**l**). Data are means ± s.e.m. Results were assessed by one-way analysis of variance followed by Tukey’s *post hoc* tests (superscript letters for factor levels *P*<0.05) on log-transformed data.

### TMA treatment mimics choline supplementation

To further confirm whether TMA mediates the beneficial effects of choline supplementation, we chronically treated LC-HFD-fed C57BL/6 mice with TMA for 6 weeks using subcutaneous osmotic minipumps and assessed their immunometabolism. We confirmed that chronic TMA treatment at 0.01 mM in a LC-HFD did not affect BW gain (**Extended Data Fig. 4g**) but effectively normalised glucose homeostasis and the Matsuda index, a readout of insulin sensitivity (**Fig. 3f-h, Extended Data Fig. 4h**). TMA treatment also improved the pro-inflammatory response to HFD-feeding and hepatic expression of acute phase proteins *Saa1*, *Saa2* and *Saa3* (**Fig. 3i-l**). These results suggest that chronic TMA treatment decouples BW gain and adiposity from low-grade inflammation and glucose homeostasis, thereby mimicking the immune and metabolic benefits observed in choline supplementation.

### TMA is a IRAK4 kinase inhibitor

To identify a direct mechanism linking TMA to metabolic and immune benefits in HFD-fed mice, we sought to identify its host pharmacological targets. The kinome, made of 518 kinases encoded the human genome^12^, represents a repertoire of critical signal transduction switches for metabolic homeostasis and inflammation. To discover specific physical interactions, we screened TMA against a panel of 456 clinically-relevant human kinases using a high-throughput assay^35,36^ (see *Methods*) and identified five preliminary hits for TMA (**Fig. 4a** and **Supplementary Table 5**). We then generated multiple-dose binding curves between TMA and each hit and confirmed that TMA physically binds IRAK4 (dissociation constant *K_d_* = 14 nM, **Extended Data Fig. 5a**) but not the other four preliminary hits (flat dissociation curves with no *K_d_* fit in **Extended Data Fig. 5b-e**, suggesting no physical binding was observed). Since *K_d_* only addresses a physical interaction in its simplest form (*i.e.:* binding), we functionally tested TMA as an IRAK4 inhibitor, by quantifying IRAK4 kinase activity in presence of ATP and increasing doses of TMA (see *Methods*) to derive a half-maximal relative inhibitory constant (*IC_50_* = 3.4 µM, **Fig. 4b**). We also confirmed that neither choline, TMAO nor DMB inhibit IRAK4 kinase activity (**Extended Data Fig. 5f-h**). We finally confirmed that TMA does not inhibit IRAK-1 either (**Extended Data Fig. 5i**), as suggested by the phenotypic convergence of *Irak-1*^-/-^ mice fed a HFD which were shown to have metabolic improvements similar to TMA treatment^37^. Altogether, this screen shows that IRAK4 is the specific kinase target for TMA, which we next validated using a range of *in vitro* and *in vivo* experiments.

**Figure 4.**
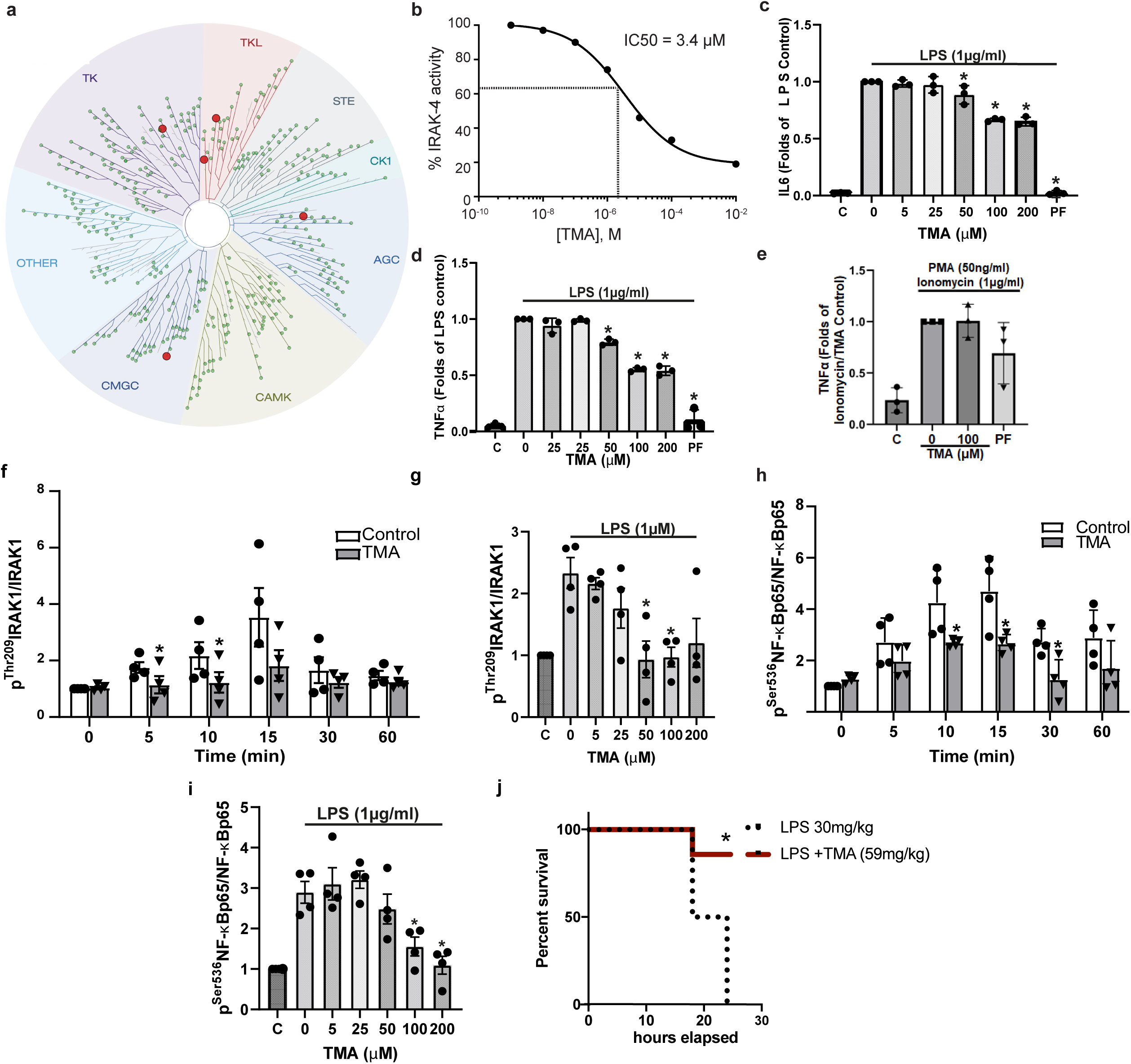
TMA inhibits IRAK4 and rescues LPS-induced TLR4-mediated pro-inflammatory response. (**a**) Kinome screen for a single dose of TMA (0.1 mM) against 456 kinases. (**b**) Functional characterization of the inhibition of IRAK4 by TMA. IRAK4 phosphorylation activity was determined in presence of various concentration of TMA and resulted in an IC_50_ of 3.4 µM. TMA preincubation (100 µM, for 30 min) suppresses LPS-induced (1 µg/ml, 4h) IL6 (**c**) and TNFα (**d**) release by human PBMCs. The values of the unstimulated controls were arbitrarily set to 1. **\***<0.05 vs LPS-stimulated control. (**e**) TMA (100μM) or the IRAK4 specific inhibitor PF06650833 (PF; 50nM) do not inhibit TNFα secretion from human PBMCs after 4h stimulation with PMA (50ng/ml) and ionomycin (1μg/ml). (**f**) Impact of TMA pre-treatment (100 µM, 30 min) on human PBMCs’ ^Thr209^IRAK1/IRAK1 phosporylation upon LPS (1 µg/ml) stimulation for the indicated times. (**g**) Human PBMCs were pre-treated (for 30 min) with the indicated concentrations of TMA and ^Thr209^IRAK1/IRAK1 phosporylation was assessed after 10 min of LPS (1 μg/ml) challenge. The time-(**h**) and dose-dependent (**i**) effect of TMA on phosphorylation of ^Ser536^NF-κBp65/NF-κBp65 of human PBMCs stimulated with LPS (1 μg/ml) was assessed as in (**f**) and (**g**). For (**f-i**), the control value was arbitrarily set to one and statistical significance (*<0.05) was assessed with unpaired Student’s *t* test. (**j**) 24-hour survival curve of mice challenged with a lethal dose of LPS (30 mg/kg) that received a single dose of TMA (59mg/kg, red line), or vehicle (black line). The kinases covered by the kinome scan assay are visualised using a phylogenetic layout. Preliminary positive hits with potential binding >35 % vs control (DMSO) are represented by a red dot whilst non-binding kinases are represented by a green dot. Kinase group names: TK, Tyrosine Kinases; TKL, Tyrosine Kinase-like; STE, STE kinase group; CK1, Cell Kinase 1; AGC, Protein kinase A, G and C families; CAMK, Calcium and Calmodulin-regulated kinases; CMGC, CMGC kinase group. Data are means ± s.e.m.

### TMA inhibits IRAK4 signalling in human PBMCs and primary human hepatocytes

IRAK4 is a central kinase in the TLR pathway sensing bacterial invasion and promoting a pyretic pro-inflammatory response in infectious contexts^38,39^. To further corroborate the inhibitory action of TMA on IRAK4 on innate immunity, we studied its effect on human peripheral blood mononuclear cells (PBMC) cytokine secretion and NF-κB signalling in response to LPS stimulation, a process exclusively mediated by IRAK4 downstream of TLR4 as originally demonstrated by Suzuki et al.^38^. Consistent with this, pre-treatment with TMA for 30 min dose-dependently blunted IL6 and TNFα secretion by PBMCs challenged with LPS after 4 hours (**Fig. 4c, d**). Moreover, IL6 and TNFα secretion was virtually abolished by pre-treatment by the IRAK4 specific inhibitor PF06650833^40^ as an additional control (100 nM). Conversely, TMA (100 µM) or the IRAK4 specific inhibitor PF06650833 (100 nM) had no significant impact on TNFα secretion by PBMCs stimulated by phorbol-ester/Ionomycin (**Fig. 4e**), similarly to human B-cells harbouring IRAK4-inactivating mutations^41^. Taken together, these findings further support our assertion that TMA acts through IRAK4 and further validate the functional inhibition of IRAK4 by TMA in innate immune cells. In addition, TMA (0.1 mM) significantly ameliorated IRAK1 phosphorylation at a site (Thr209) phosphorylated by IRAK4^42^ upon short-term stimulation of PBMCs with LPS (up to 60min, **Fig. 4f**). TMA also inhibited ^Thr209^IRAK1 phosphorylation dose-dependently after LPS stimulation of PBMCs (**Fig. 4g**). Similarly, TMA time- and dose-dependently suppressed NF-κBp65 phosphorylation in LPS-challenged PBMCs (**Fig. 4h, i**). In agreement with the cellular PBMC experiments a single TMA dose significantly improved mouse survival in a 24-hour LPS septic shock experiment (**Fig. 4j**), which shows similar protection to the Irak4 kinasedead mice^43^.

We then sought to test IRAK4 inhibition in a HFD context, the main free fatty acid (FFA) in our HFD being palmitate, a saturated FFA triggering TLR4 signalling. We therefore used a primary human hepatocyte model of low-grade inflammation and insulin resistance^44^. In basal conditions (0 min), TMA (0.1 mM, 30 min) pre-treatment resulted in a significant decrease in ^pThr345/Ser346^IRAK4/IRAK4, ^pThr209^IRAK1/IRAK1 and ^pSer176/180^IKKαβ/IKKβ. PA (200 μM, 60 min) administration resulted in increased ^pThr209^IRAK1/IRAK1, ^pSer176/180^IKKαβ/IKKβ and ^pSer536^NF-κBp65/NF-κBp65 (**Extended Data Fig. 6a-d**), which were normalized by TMA treatment. The TMA effect was significant in a 2-way ANOVA for ^pThr345/Ser346^IRAK4/IRAK4 (p=0.04), ^pThr209^IRAK1/IRAK1 (p=0.03) and ^pSer176/180^IKKαβ/IKKβ (p<0.001). The normalization of the phosphorylation ratios in the TLR4 pathway by TMA resulted in a normalization of PA-induced IL-6 secretion in the cell medium after 4 hours (**Extended Data Fig. 6e**). TMA also prevented the negative impact of PA (200 μM, 4 h) on insulin action (specifically on insulin-induced ^pSer473^Akt1/Akt1), indicating a beneficial TMA effect on insulin signaling (**Extended Data Fig. 6f**). TMA also tended to decrease ^pThr183/Tyr185^SAPK/JNK/SAPK/JNK induced by PA (200 μM, 60 min, **Extended Data Fig. 5j**) but had no significant effect on p38MAPK (**Extended Data Fig. 5k**). Collectively, our results demonstrate that TMA is an IRAK4 kinase inhibitor ameliorating LPS inflammatory signalling in PBMCs and mice and normalising palmitate-induced low-grade inflammation and aberrant insulin signalling *in vitro*, requiring further assessment in *Irak4*^-/-^ mice.

### *Irak4^-/-^* mice are protected against HFD-induced immune and metabolic dysregulations

IRAK4 is a key kinase required for defence against pyogenic infections in acute contexts^38,39^. To further test whether this kinase plays a role in HFD-induced chronic low-grade inflammation and glucose homeostasis, we fed 5-week-old *Irak4^-/-^* mice^45^ and wild-type (WT) littermates in C57BL/6 background (**Extended Data Fig. 7a** for genotyping) a LC-HFD, to avoid potential confounding effects from TMA for 8 weeks before assessing circulating cytokines, expression of hepatic inflammatory genes and acute-phase markers and metabolic homeostasis (**Fig. 5a-h**). Irak4^-/-^ mice presented improved glycemic control compared to wild-type littermates (**Fig.5 a,b**). Likewise, the inflammatory response to LC-HFD observed in WT was obliterated in *Irak4^-/-^* littermates (**Fig. 5c-g**). There was a similar trend for *Saa3* (**Fig. 5h**), whilst there was no effect on BW gains (**Extended Data Fig. 7b**). Hence, similarly to TMA treatment, genetic ablation of IRAK4 abolishes the HFD-induced pro-inflammatory response and IGT thereby decoupling obesity from impaired glucose tolerance and low-grade inflammation with a similar phenotype to *Irak1* deficiency^37^.

**Figure 5.**
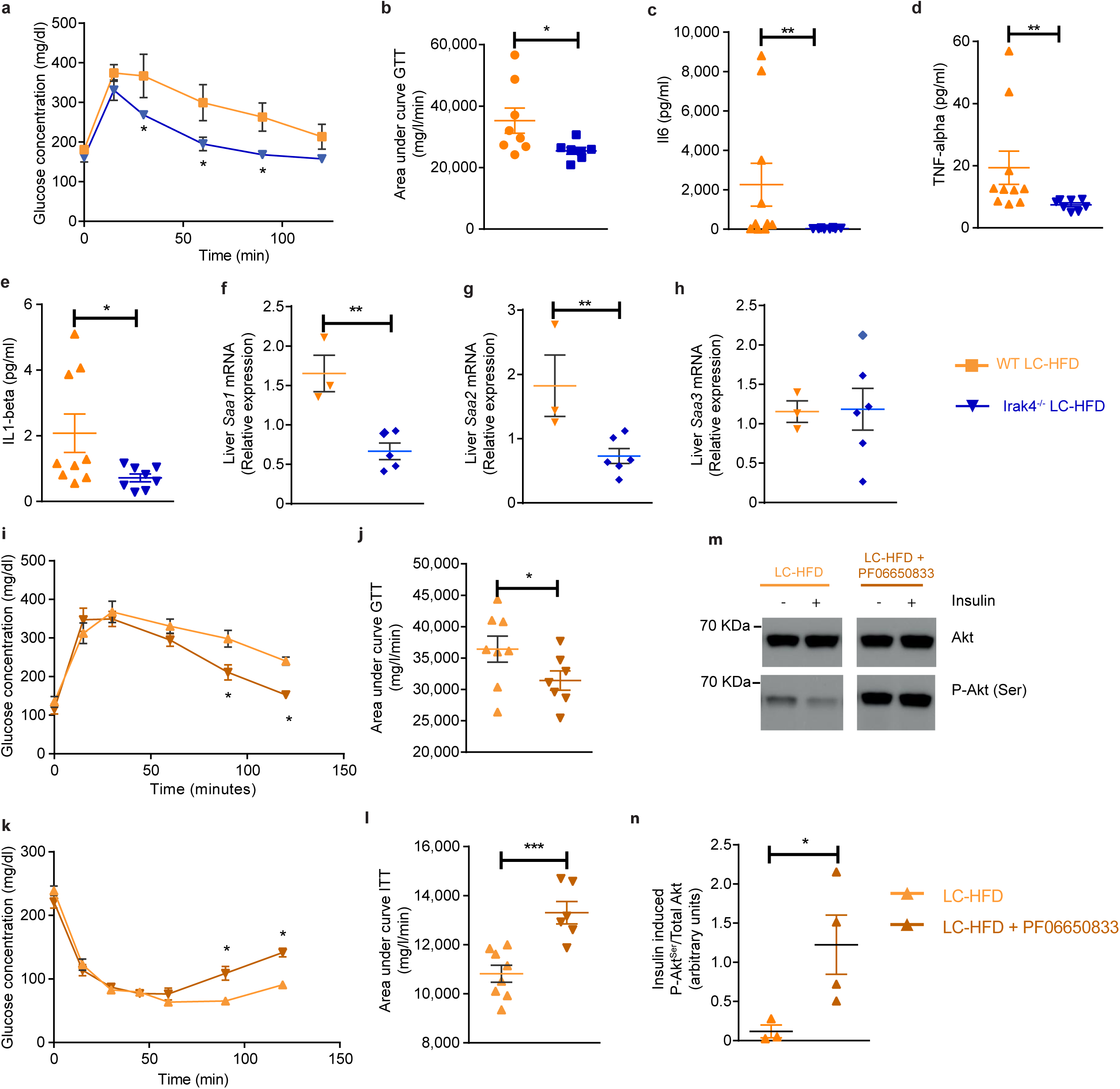
IRAK4 deficiency corrects IGT and pro-inflammatory response after 8 weeks of low choline HFD feeding in C57BL/6 mice and a chemical inhibitor of IRAK4 mimics the effects on glucose homeostasis. (**a-h**) 5-week-old *Irak4*^-/-^ mice and wild-type (WT) littermates were fed a low-choline HFD (10 WT HFD and 8 *Irak4*^-/-^ HFD) and were phenotyped after 8 weeks. (**a**) Plasma glucose concentration during an ipGTT. (**b**) Area under the curve of the plasma glucose concentration during an ipGTT. (**c-e**) Circulating cytokines (**c**) IL6, (**d**) TNFα, (**e**) IL1β. (**f-h**) Liver mRNA expressions for *Saa1* (**f**), *Saa2* (**g**) and *Saa3* (**h**) genes. (**i-n**) Mice were weaned at 3 weeks and fed a low-choline HFD before being fitted with osmotic minipumps delivering a chronic circulating dose of PF06650833 (50 nM) for 6 weeks. (**i**) Plasma glucose concentration during an ipGTT. (**j**) Area under the curve of the plasma glucose concentration during an ipGTT. (**k**) Plasma glucose concentration during an ipITT. (**l**) Area under the curve of the plasma glucose concentration during an ipITT. (**m-n**) Western-blot analysis of liver Akt phosphorylation. Data are means ± s.e.m. Student’s *t* test (*P*<0.05) on log-transformed data.

### Pharmacological inhibition of Irak4 normalises glucose metabolism

Since the *Irak4^-/-^* mice lack the whole protein, we compared the knock-out phenotype with the phenotype of PF06650833, a recently discovered chemical inhibitor of the human IRAK4 protein^40^ with promising results in a phase-I trial for rheumatoid arthritis^46^. Treatment with PF06650833 improved BW in LC-HFD mice (**Extended Data Fig. 7c**). This inhibitor also yielded significant improvements in plasma glycemia at the latter time-points of the GTT and insulin tolerance test (ITT) and in cumulative glycemia (**Fig. 5i-l**), which was mirrored by an increase in Akt phosphorylation (**Fig. 5m-n**). These results collectively show that specific chemical inhibition of IRAK4 leads to significant improvements in glycemic control, insulin sensitivity and insulin signalling. Further, they suggest that IRAK4 could constitute a clinically relevant target in IR and related disorders.

## DISCUSSION

The discovery that TMA is a kinase inhibitor controlling IRAK4, a central kinase involved in innate immunity, is a major finding that provides an attractive mechanism for the metabolic and immune improvements observed with choline-supplementation in HFD contexts (**Fig. 6**).

**Fig. 6.**
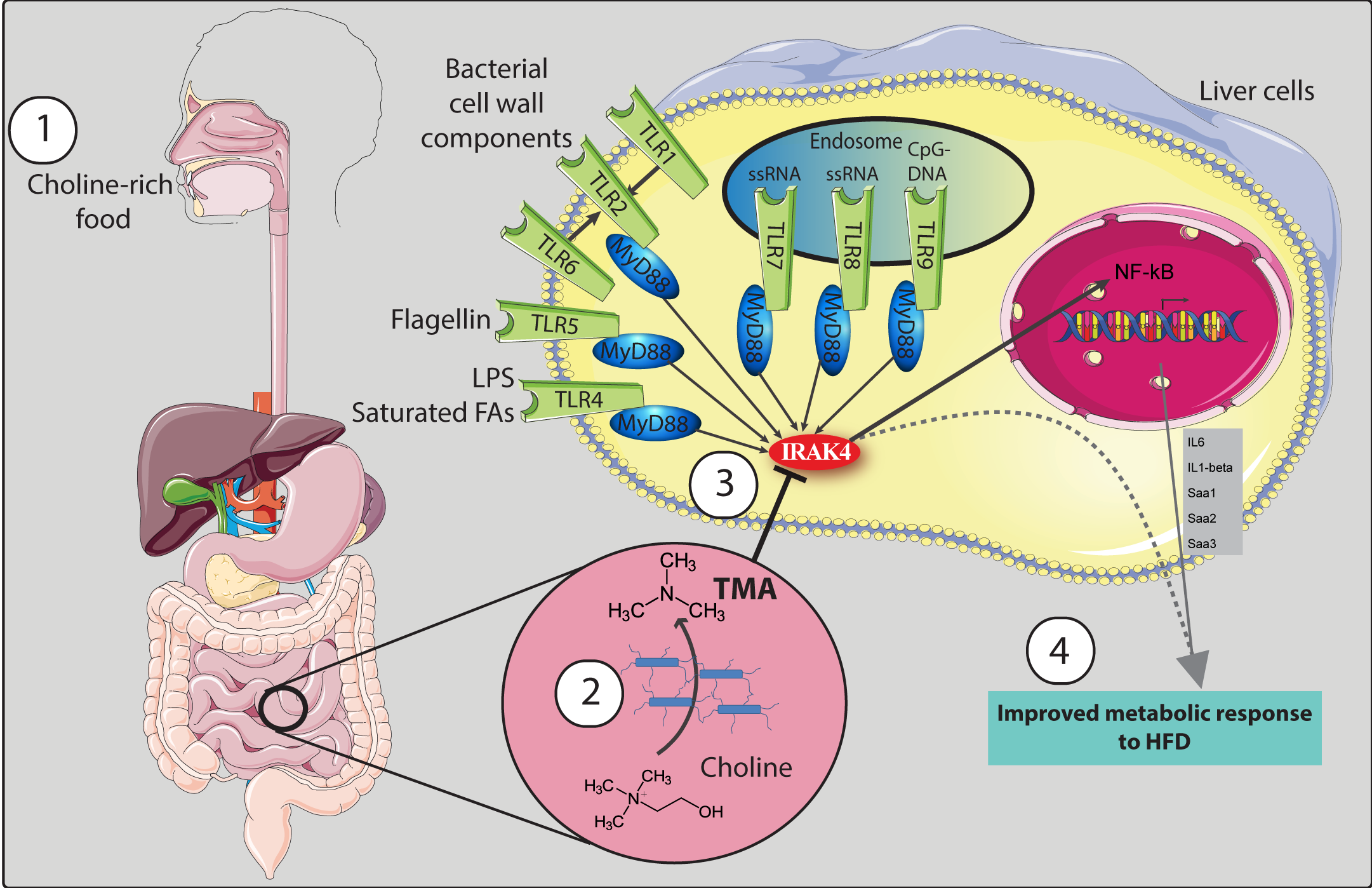
Overview of effects of IRAK4 inhibition by TMA on metabolic response to HFD. TMA is synthesized from dietary choline (1) by the gut microbiota (2) and specifically inhibits IRAK4 activity (3). This inhibition blunts the TLR signalling pathway (induced by LPS and saturated free fatty acids) leading to an improvement of metabolic response to HFD (e.g. glucose homeostasis).

IRAK4 is the first regulatory checkpoint downstream of MyD88, the adapter protein connecting IRAK4 to at least six TLRs sensing bacterial compounds/components^47^. IRAK4 deficiency is associated with bacterial infections in humans^39^ and in mice^48^. Consistent with the involvement of the TLR signalling pathway at the crossroads between gut microbiota and dietary lipids^49,50^, IRAK4 deletion and its inhibition by TMA and PF06650833^40^ rescued HFD-induced low-grade inflammation and IR^3,51,52^, highlighting new and unexpected roles for this microbial metabolite and its target kinase in immunometabolism. The relative IC_50_ (3.4 µM) we determined is two times smaller than the plasma isotopic quantifications obtained by UPLC-MS/MS in the HFD group supplemented with choline. These quantifications are comparable to circulating TMA levels previously reported in normal human plasma, ranging between 0.42 and 48 µM^53,54^, which makes this IC_50_ particularly relevant with respect to pathophysiological mechanisms: dosing mice with 0.01 mM TMA (3x IC_50_) was sufficient to improve inflammatory and metabolic responses. TMA can therefore be considered a “microbial signalling metabolite”^55^ sending a negative feedback signal to a pathway triggered by influx of saturated free fatty acids in HFD contexts, this mechanism participating in maintaining a low immunological footprint and improved metabolic homeostasis in a symbiotic relationship. TMA’s role as an IRAK4 inhibitor could, for instance, explain some of the beneficial effects reported for choline supplementation in NAFLD patients^56^ and of improved IR observed in those consuming higher choline diets in a healthy genetically uniform human population^57^. Altogether, our results on IRAK4 inhibition and ablation provide further insight in the phenotypic convergence between *Myd88*, *Irak4* and *Irak1* knock-out mouse models^37,58^, suggesting that the gut microbiota, through TMA, proceeds with a targeted “hijacking” of the TLR signal transduction machinery to the host’s benefit resulting in metabolic and immune improvements.

Our finding that TMA improves metabolic health in the context of HFD is surprising given the extensive evidence linking higher levels of circulating TMAO (after phase-1 *N*-oxidation of TMA in the liver) with worst cardiovascular outcomes, particularly in patients with established atherosclerosis and thrombosis^14^. Our results show that the TMA-lyase inhibitor DMB blocks TMA production and abrogates the metabolic benefits brought by choline supplementation in the HC-HFD mice, suggesting that the benefits come from conversion of choline to TMA. Our findings are supported by the discovery that in diabetic mice knockdown of the liver gene (*fmo3*) that converts TMA to TMAO, an intervention that would increase TMA levels, improves IR and attenuates atherosclerotic burden^23^. Similarly, in an epidemiological study in a genetically homogeneous population, higher choline consumption associated with improved insulin sensitivity^57^.

Our work is consistent with the idea that TMA and TMAO may well play contrasting roles that are mechanism- and context-dependent. To illustrate context dependence, TMAO, which has been associated with poor cardiovascular outcomes in humans also beneficially reduces blood-brain barrier permeability^59^, independently of its role in CVD. To illustrate the target-dependence, TMAO augments Ca^2+^-mediated platelet aggregation after thrombin treatment. TMAO also binds to and activates purified Protein Kinase RNA-Like ER Kinase (PERK) whilst structurally related metabolites such as choline do not^60^. Similarly, we discovered that TMA binds to and inhibits IRAK4 whilst TMAO does not (ED figure 5g), suggesting an unexpected and independent mechanism of action. Our observations on IRAK4 inhibition by TMA and in *Irak4* knock-out mice are consistent with observations in Myd88 and Irak1 knock-out mice, resulting in improved glucose tolerance and insulin sensitivity. Consistent with our observations, *Fmo3* deletion, which as a result prevents *N*-oxidation of TMA into TMAO, improved IR in mice^23^. Therefore, we submit that our study reveals divergent context-dependent actions between TMA and its phase-II metabolite TMAO in an HFD context in a “yin-yang relationship”.

In conclusion, through the combination of *ex vivo*, *in vitro* and *in vivo* approaches, we reveal a unique mechanism in which TMA acts as a gut microbial signalling metabolite inhibiting IRAK4, a central molecular target in the TLR pathway, thereby allowing the gut microbiota to control HFD-induced pro-inflammatory response(s) and IR. The kinome represents a key repertoire of regulatory host targets for microbial signals and the uncharted microbiome– kinome crosstalk requires further investigation. Our work can guide human trials where the efficacy of IRAK4 inhibitors available for human use or dietary interventions that increase TMA bioavailability can be evaluated in the context of obesity and IR. Moreover, since IR is an independent risk factor of CVD in humans^61^, our work uncovers a novel strategy to alleviate obesity-associated increased cardiovascular risk. By highlighting the physiological and therapeutic roles of TMA and IRAK4 on HFD-induced low-grade inflammation and IR, we anticipate that its immunomodulatory properties extend beyond IR to a wider range of human pathologies involving TLR signalling and modulation of innate immunity.

## Acknowledgments

This work was supported by grants from the Wellcome Trust (Functional Genomics Initiative grant Biological Atlas of Insulin Resistance 06678 to JKN, DG and JS) and the European Commission (METACARDIS HEALTH-F4-2012-305312 to DG and MED, Institut Mérieux through a grant awarded to MED, Spanish Institute of Health Research (FIS) grant (PI18/01022 and PI21/01361), by Fondo Europeo de Desarrollo Regional (FEDER) funds, the Catalan Government (AGAUR, #SGR2017-00734, ICREA Academia Award 2021) to JMFR and Canadian Institutes of Health Research (CIHR) grants (MOP-49413 & MOP-142471) awarded to PPL. The laboratory of M.E.D. has received funding by the UK Medical Research Council (MRC grants “μNeuroInf” MR/M501797/1 and “National Mouse Genetics Network Microbiome Cluster” MR/W022532/1), and by grants from the French National Research Agency (ANR-10-LABX-46 [European Genomics Institute for Diabetes]), from the National Center for Precision Diabetic Medicine – PreciDIAB, which is jointly supported by the French National Agency for Research (ANR-18-IBHU-0001), by the European Union (FEDER), by the Hauts-de-France Regional Council (Agreement 20001891/NP0025517) and by the European Metropolis of Lille (MEL, Agreement 2019_ESR_11) and by Isite ULNE (R-002-20-TALENT-DUMAS), also jointly funded by ANR (ANR-16-IDEX-0004-ULNE), the Hauts-de-France Regional Council (20002845) and by the European Metropolis of Lille (MEL). This research was also supported by the National Institute for Health Research (NIHR) Imperial Biomedical Research Centre (MED, S.D.G.). DG held a Wellcome Trust Senior Fellowship in Basic Biomedical Science (057733). JEF was supported by the Medical Research Council (grant number MR/K020919/1). PA is the recipient of a Career Development Award from the Medical Research Council (Grant No. MR/Y010051/1). PDC is a senior research associate and AE a research associate at FRS-FNRS (Fonds de la Recherche Scientifique). PDC is the recipient of grants from FNRS. LH was in receipt of an MRC Intermediate Research Fellowship in Data Science (grant number MR/L01632X/1, UK Med-Bio). This work was supported by FRFS-WELBIO under grant: WELBIO-CGR-2017, by the Funds InBev-Baillet Latour (Grant for Medical Research 2015) and ERC Starting Grant 2013 (European Research Council, Starting grant 336452-ENIGMO).

## Author Contributions

JC, PA, BAS and SG performed cell-based assays, JC performed multivariate statistics and interpreted the results. JC, FB, JFF, AE, HP, LZ, DS, ARR, PDC, VC, RM, SC and CG contributed *in vivo* experiments, RHB, AM, ALN, LMG, SG, JMMN, JL, CLB and SP performed various experiments. LH analysed microarray data. JS, DG, JKN, PPL, PDC, JMFR, NJG and MED conceived the project, mentored and supervised its participants and interpreted its results. MED performed metabolic profiling experiments, analysed data and interpreted the results. JC, PA, NJG, DG, PPL, PDC, and MED wrote the manuscript. All authors edited and approved the manuscript.

## Online Content

Methods, along with any additional Supplementary display items and Source Data, are available in the online version of the paper; references unique to these sections appear only in the online paper.

## Online Methods

### Protocols

All experimental procedures involving mice were carried out in accordance with U.K. Home Office, Canadian Council on Animal Care, the ethics committee of the French Research Ministry (authorization number 00486.01), Belgian Law of May 29, 2013 regarding the protection of laboratory animals (agreement number LA1230314) and local guidelines on animal welfare and license conditions and the University of Oxford, University of Ottawa, Université Pierre et Marie Curie and Université catholique de Louvain guidelines on animal welfare.

### PBMC isolation and cell-based assays

PBMCs were isolated as previously described^62^ following local ethical approval from Imperial College Research Ethics Committee (19IC5372). Briefly, human peripheral blood samples (32 ml) were collected from donors in BD Vacutainer Cell Preparation Tubes containing Sodium Heparin/Ficoll (BD Biosciences, UK). PBMCs were separated by centrifugation at 1600 ***g*** for 30 min at room temperature, followed by three washing steps with PBS and 10% FBS (LabTech, UK), and were centrifuged at 520 RCF for 10 min after each wash. Donors were females aged between 25 and 32 years. All subjects were healthy volunteers by self-declaration and provided written informed consent.

### LPS stimulation and IL6 and TNFα release quantification

Human PBMCs (1x10^5^ per condition in duplicate), freshly isolated as described above, were serum-starved in 0.1% v/v FCS RPMI-1640 (Sigma Cat. R7638) in the presence of TMA (Sigma Cat.No. 72761) or the IRAK4 inhibitor PF06650833 (0.5μM; Tocris Cat.No.6373; PF thereafter), as indicated in the corresponding Figures, for 30 min. PBMCs were subsequently stimulated with 1 μg/ml LPS (Sigma Cat.No. L2630) for 4 h (37 °C, 5% CO_2_). For the PMA/Ionomycin stimulation experiments, human PBMCs isolated as above (1x10^5^ per condition) pre-incubated with TMAO (100 μM), PF06650833 (0.5μM) or vehicle as indicated for 30min were then stimulated with 50ng/ml Phorbol 12-myristate 13-acetate (PMA; Sigma Cat.No. 79346) and 1μg/ml ionomycin (Sigma Cat.No. I0634) for 4h. Next cells were pelleted by centrifugation (200 ***g*** for 10 min) and supernatants were collected and stored at -20 °C until further use. IL6 and TNFα were measured in the media diluted 1:10 in RPMI-1640 using the DuoSet ELISA kits from R&D (Cat.No. DY206 and Cat.No. DY210 from R&D, respectively) according to the manufacturer’s instructions. For PMA/ionomycin TNFα measurements the media were un-diluted. For all treatments *N*=4 biological repeats.

### PBMCs IRAK1 and NF-κBp65 phosphorylation in response to LPS

Freshly isolated human PBMCs (1x10^6^/ml per condition) were added to 96-well plates coated with *L*-poly-lysine for 30 min (100 ng/ml, Sigma Cat.No. P4707) in RPMI-1640 containing 20% FCS and were left to attach to the wells overnight at 37 °C, 5% CO_2_. The following day media were discarded and cells were incubated in serum-free RPMI-1640 containing 100 μM TMA, or RPMI-1640 vehicle for 30 min. PBMCs were then stimulated with LPS (1 μg/ml) for the indicated times in corresponding Figure legends, for up to 60 min. At the end of the stimulation PBMCs were fixed with 8% paraformaldehyde (Sigma Cat.No. F8775) at room temperature for 20 min. Phosphorylation of IRAK1 (at Thr209) or NF-κBp65 (at Ser536) was determined in fixed PBMCs using commercially available ELISA kits (Cat.No.LS-F1401-1 or Cat.No.LS-F891-1 respectively, both from LSBio) according to the manufacturer’s instructions. Readings were normalized to the levels of total-IRAK1 protein determined in sister wells treated identically. The dose-dependent impact of TMA (see corresponding Figures) on phospho levels of IRAK1 or NF-κBp65 upon LPS (1 µg/ml) stimulation of PBMCs (for 10 min or 15 min, respectively) was also determined using the commercially available kits from LSBio as described above.

### Cell-based assays in primary human hepatocytes

Cryopreserved primary human hepatocytes (HH) were commercially sourced (Innoprot, Bizkaia, Spain) and cultured with hepatocytes medium (Innoprot) supplemented with 5% fetal bovine serum, 1% hepatocytes growth supplement (mixture of growth factors, hormones and proteins necessary for culture of primary hepatocytes) and 100 U/mL penicillin and streptomycin. HH were grown on poly-*L*-lysine pre-coated cell dishes at 37 °C and 5% CO_2_ atmosphere following manufacturer’s recommendations. The following experiments were performed 24 h after seeding: i) First, palmitic acid (PA, 200 μM) administration (0 and 60 min) with or without trimethylamine hydrochloride (TMA, 0.1 mM, 30 min) pre-treatment plus co-administration during PA exposure were assayed. PA solution was prepared as previously reported^44,63^. In this experiment, the effect of PA and TMA on IRAK1, IRAK4, NF-κBp65, IKKαβ, SAPK/JNK and p38MAPK activity was analyzed. ii) In a second experiment, the effect of TMA (0.1 mM, 30 min) pre-treatment plus co-administration during vehicle or PA exposure for 4 h on insulin action and on IL6 release were tested. Insulin action was evaluated measuring ^pSer473^Akt1/Akt1 after insulin (100 nM, 10 min) stimuli.

^pThr209^IRAK1, total IRAK1, ^pSer536^NF-κBp65, total NF-κBp65, ^pSer473^AKT1, total AKT1 and GAPDH (as an endogenous control) were measured with specific colorimetric Cell-Based ELISA Kits [CytoGlow^TM^ IRAK1 (catalog n°: CBP1425), CytoGlow^TM^ NF-κBp65 (catalog n°: CBP1633) and CytoGlow^TM^ AKT1 (catalog n°: CBP1490), Assay Biotechnology Company, Inc, CA, USA] following manufacturer’s instructions. For data analysis, optical density of target proteins (read at 450 nm) was normalized with cell nuclei crystal violet staining (read at 595 nm), which was proportional to cell counts. The analysis of cell-based assays was performed in a blind manner.

_pThr345/Ser346IRAK4/IRAK4, pSer176/180IKKαβ/IKKβ, pThr183/Tyr185SAPK/JNK and pThr180/Tyr182_p38MAPK/p38MAPK were determined by western blot. Briefly, hepatocyte proteins were directly extracted in radioimmnuno precipitation assay (RIPA) buffer (0.1% SDS, 0.5% sodium deoxycholate, 1% Nonidet P-40, 150 mM NaCl, and 50 mM Tris-HCl, pH 8.0), supplemented with protease inhibitors (1 mM phenylmethylsulfonyl fluoride). Cellular debris and lipids were eliminated by centrifugation of the solubilized samples at 13000 rpm for 10 min at 4 °C, recovering the soluble fraction. Protein concentration was determined using the RC/DC Protein Assay (Bio-Rad Laboratories, Hercules, CA). RIPA protein extracts (20 μg) were separated by SDS-PAGE and transferred to nitrocellulose membranes by conventional procedures. Membranes were immunoblotted with antibodies against the following proteins: ^pThr345/Ser346^IRAK4 (#11927), IRAK4 (#4363), ^pSer176/180^IKKαβ (#2694), IKKβ (#8943), ^pThr183/Tyr185^SAPK/JNK (4668), SAPK/JNK (#9258), ^pThr180/Tyr182^p38MAPK (#9215), p38MAPK (#9212), all purchased from Cell Signaling Technology, Inc (CST, MA, USA) and β-actin (sc-47778, Santa Cruz Biotechnology, CA, USA). Anti-rabbit IgG and anti-mouse IgG coupled to horseradish peroxidase was used as a secondary antibody. Horseradish peroxidase activity was detected by chemiluminescence, and quantification of protein expression was performed using scion image software. IL6 concentration in hepatocyte media was measured using Human IL-6 Quantikine ELISA Kit (D6050, R&D Systems, Inc, MN, USA). All these experiments were performed at least in three sample replicates.

### Mouse Models

#### Longitudinal HFD-feeding in mice

All experiments were approved by the ethical committee of the University of Oxford. Male mice from C57BL/6J inbred strain were bred in our animal facility by using a stock originating from The Jackson Laboratory. At 5 weeks of age, groups of n=8-10 mice were transferred to a 40 % w/w HFD (65% kcal) (Special Diets Services), containing 32 % lard and 8 % corn oil, whereas control groups remained on a 5 % Low Fat Diet (CHD) (B & K Universal) for up to 6 months. Detailed diet formulations were published previously^13^ and summarised in **Supplementary Table 6**.

Mice were housed under a 12 h–12 h light–dark cycle. For physiological profiling, several mouse groups fed CHD or HFD were tested to assess consistency of results and discard any impact of potential batch effects. Intra peritoneal glucose tolerance tests (ipGTTs) were performed in mice after 2-, 3-, 5-and 7-month-old mice after an overnight fast, as previously published^64^ (see also metabolic phenotyping below). Four days after the GTT, 24 h urinary samples (9 a.m. to 9 a.m.) were collected from mice maintained in individual metabolic cages. Urinary samples collected in a solution of 1 % (wt/vol) sodium azide were centrifuged to remove solid particles and kept at -80 °C until assayed. After an overnight fast, mice were killed by exsanguination. Plasma was separated by centrifugation and stored at -80 °C until ^1^H-NMR analysis.

#### Choline supplementation on HFD

At 5 weeks of age, mice were fed either a chow diet containing 2 g of choline/kg of diet (Research Diets, D12450J), a low-choline HFD containing 2 g of choline/kg of diet (Research diets, D12492), a high-choline HFD containing 17 g of choline/kg of diet (Research diets, D16100401), a HC-HFD containing 1% of DMB or a HC-HFD combined with a cocktail of antibiotics (0.5 g/L vancomycin hydrochloride, 1 g/L neomycin trisulfate, 1 g/L metronidazole, 1 g/L ampicillin sodium) in drinking bottles (n=6-10 per group) for 8 weeks (see diet formulations in **Supplementary Table 7**). Mice then were killed by decapitation and organs were dissected and weighed.

#### Irak4^-/-^ mice

*Irak4^-/-^* mice on C57BL/6J background as already described^38,45^ were bred with C57BL/6J mice (The Jackson Laboratory), and the F1 offspring were subsequently bred to produce *Irak4^-/-^* mice and WT littermate used for this study. Mice were breaded and genotyped at animal facility of University of Ottawa Heart Institute. The following primers were used for genotyping. Irak4 KO 5’-tga atg gaa gga ttg gag cta cgg ggg t -3’; Irak4 common 5’- gaa cac gct ccc agg tct ctt tcc aac; and Irak4 WT 5’- tct tct acc tga aat atg aaa gat tcc t -3’. The PCR reaction were run at 94 °C 60s, 60 C 60s, 72 C 60s for 40 cycles. 10-12 weeks old mice were feed with HFD for eight weeks and were killed by decapitation and organ were dissected and weighed at the end of the study.

#### Chronic TMA and PF06650833 treatment in LC-HFD-fed mice

Five-week-old C57BL/6J mice (Charles River) were housed a week before experiment in a controlled environment. Mice were housed under a 12 h–12 h light–dark cycle. At day 0, the 10-week-old mice were anaesthetised with isoflurane (ForeneH, Abbott). Mini-osmotic pumps were implanted subcutaneously (Model 2006, Alzet) (flow rate: 0.15 mL/h, total filling volume: 200 mL, delivery duration: 42 days) as previously described^65^. The osmotic mini-pump contains either vehicle or TMA (0.1 mM in circulation) or PF06650833 (50 nM in circulation). After six weeks of metabolite treatment, mice were killed by decapitation and organs were dissected and weighed.

#### Septic shock mouse model

Six-week-old male mice in a C57BL/6J background were purchased from Charles River (Calco, Italy) and maintained in a controlled environment for 2 weeks to acclimate them to local conditions. Animal experiment protocol was approved by local and national committees in charge (Tor Vergata University Institutional Animal Care and Use Committee and Ministry of Health, license no. 265/2019-PR) and conducted in accordance with accepted standard of humane animal care. Mice were intraperitoneally injected with 59mg/kg TMA (Sigma Cat.No. 72761) (treatment group, n=7), or PBS alone (control group, n=6) 30 minutes before LPS injection [30 mg/kg of LPS (Sigma Cat.No. L2630) in sterile PBS by intraperitoneal injection]. The survival of the mice was monitored every 4 h for 36 h.

### Physiological phenotyping

After 4 weeks of treatment an ipGTT test (2 g/kg) was performed in conscious mice following an overnight fast. Blood was collected from the tail vein before glucose injection and 30, 60, 90 and 120 min afterwards. Blood glucose levels were determined using an Accu-Check® Performa (Roche Diagnostics, Meylan, France). Additional blood samples were collected at baseline and 30 min after glucose injection in Microvette® CB 300 Lithium Heparin (Sarstedt, Marnay, France). Plasma was separated by centrifugation and stored at - 80 °C until insulin radioimmunoassay. Circulating insulin levels were determined using Insulin ELISA kits (Mercodia, Uppsala, Sweden). The Matsuda insulin sensitivity index was calculated as previously published^31^.

After 5 weeks, we performed an ITT. 5-hour-fasted mice were injected intraperitoneally with insulin (0.75 mU/g, Actrapid, Novo Nordisk). Blood glucose levels were measured immediately before and 15, 30, 45, 60, 90 and 120 min after insulin injection with a standard glucose meter (Accu-Check, Roche, Basel, Switzerland) on the tip of the tail vein.

### Gene expression

Groups of 6 mice showing consistent pathophysiological profiles in response to CHD or HFD treatment were selected for microarray analysis performed as previously described^64^ and data were deposited in ArrayExpress under accession number E-MEXP-1755. The Bioconductor^66^ package Limma^67^ was used to generate the list of differentially expressed genes. Gene ontology was implemented using Enrichr^68^ and signalling pathway impact analysis using SPIA^69^.

For qPCR analysis, total RNA was prepared from tissues using TriPure reagent (Roche). Quantification and integrity analysis of total RNA were performed by analysing 1 µL of each sample in an Agilent 2100 Bioanalyzer (Agilent RNA 6000 Nano Kit, Agilent). cDNA was prepared by reverse transcription of 1 mg total RNA using a Reverse Transcription System kit (Promega). Real-time PCR was performed with the StepOnePlus real-time PCR system and software (Applied Biosystems) using Mesa Fast qPCR (Eurogentec) for detection according to the manufacturer’s instructions. RPL19 RNA was chosen as the housekeeping gene. All samples were performed in duplicate in a single 96-well reaction plate and data were analysed according to the 2_DDCT method. The identity and purity of the amplified product were assessed by melting curve analysis at the end of amplification. The primer sequences for the targeted mouse genes are: SAA1 forward CAT-TTG-TTC-ACG-AGG-CTT-TCC, SAA1 reverse GTT-TTT-CCA-GTT-AGC-TTC-CTT-CAT-GT, SAA2 forward GGG-GTC-TGG-GCT-TCC-CAT-CT, SAA2 reverse CCA-TTC-TGA-AAC-CCT-TGT-GG, SAA3 forward CGC-AGC-ACG-AGC-AGG-AT, SAA3 reverse CCA-GGA-TCA-AGA-TGC-AAA-GAA-TG as reported^70^. Q-RT-PCR assays were performed in a single batch with the personnel blinded to treatment groups.

### Circulating cytokine quantification

Circulating cytokines were quantified using MSD V-PLEX Plus Proinflammatory Panel 1 kit. Plasma samples were diluted two times in diluent provided and the experiment was processed as requested by the manufacturer and read on a SECTOR imager 2400. Cytokine assays were performed in a single batch with the personnel blinded to treatment groups.

### Western-blotting

Western blot analyses were performed as described previously^71^. To analyze the insulin signaling pathway, mice were allocated to either a saline-injected subgroup or an insulin-injected subgroup so that both subgroups were matched in terms of BW and fat mass. They then received 1 mU insulin/g BW (Actrapid; Novo Nordisk A/S, Denmark) or an equal volume of saline solution. Three minutes after injection, mice were killed and liver was harvested. 30 mg of liver were homogenized in 680 µL of RIPA buffer containing a cocktail of protease and phosphatase inhibitors. The homogenate was then centrifuged at 12,000 ***g*** for 20 min at 4 °C. Equal amounts of proteins were separated by SDS–PAGE and transferred to nitrocellulose membranes. Membranes were incubated overnight at 4 °C with antibodies diluted in Tris-buffered saline Tween-20 containing 1% bovine serum albumin: p-Akt Ser473 (1:1,000; #4060, Cell Signaling), Total Akt (1:1000, #9272S, Cell Signalling), p-NF-κB (1:3000, #ab86299, AbCam), Total NF- κB p65 (1:3000, #8242, Cell Signalling). Insulin-induced pAkt/total Akt corresponds to the ratio between pAkt/Akt in insulin treated mice and pAkt/Akt in saline treated mice. For these western blots, the membranes were stripped and re-probed with a β-actin antibody as a loading control.

### ^1^H-NMR spectroscopy and multivariate statistics

Mouse urine and plasma samples were prepared and measured on a spectrometer (Bruker) operating at 600.22 MHz ^1^H frequency; full resolution ^1^H-NMR spectra were then processed and analysed using orthogonal partial least square discriminant analysis (O-PLS-DA) as described previously^13^. Variance component analysis was performed as described previously^72^. ^1^H-NMR profiling was performed in a single batch with the personnel blinded to treatment groups. The O-PLS-DA model was validated using 10,000 random permutations of the original class membership variable to explain (i.e. diets, treatments or genotypes), as described previously^72^.

### Plasma methylamine quantification by UPLC-MS/MS

Methylamines were quantified as previously described^29,30^. Plasma samples (20 μL) were spiked with 10 μL Internal Standard (IS) solution (^13^C_3_/^15^N-TMA, d_9_-TMAO, d_4_-choline, d_3_-carnitine and d_9_-betaine in water; 1 mg/L) and 45 μL of ethyl 2-bromoacetate solution (15 g/L ethyl 2-bromoacetate, 1% NH_4_OH in acetonitrile) were added to derivatize methylamines (TMA and ^13^C_3_/^15^N-TMA) to their ethoxy-analogues, completed after 30 min at room temperature. Methylamines were derivatised with ethyl bromoacetate to increase its sensitivity (the underivatized form was giving low response to the mass spectrometer due to its low molecular weight) and enhance its chromatographic performance. 935 μL of protein/lipid precipitation solution (94% acetonitrile/5% water/1% formic acid) was added; samples were centrifuged for 20 min (4°C, 20,000 ***g***) and were transferred to UPLC-autosampler vials. Sample injections (10 μL loop) were performed on a Waters Acquity UPLC-Xevo TQ-S UPLC-MS/MS system equipped with an Acquity BEH HILIC (2.1 × 100 mm, 1.7 μm) chromatographic column. An isocratic elution was applied with 10 mM ammonium formate in 95:5 (v/v) acetronitrile:water for 14 min at 500 μL/min and 50 °C. Positive electrospray (ESI+) was used as ionization source and mass spectrometer parameters were set as follows: capillary, cone and source offset voltages at 500, 93 and 50 V, respectively, desolvation temperature at 600 °C, desolvation/cone/nebulizer gases were high-purity nitrogen at 1000 L/hr, 150 L/hr and 7 bar, respectively. Collision gas was high-purity argon. Mass spectrometer was operated in multiple reactions monitoring (MRM) mode. The monitored transitions were the following: for derivatized-TMA, +146 -> +118/59 *m*/*z* (23/27 V); for derivatised-^13^C_3_/^15^N-TMA, +150 -> +63 (27V); for TMAO, +76 -> +59/58 *m*/*z* (12/13 V); for d_9_-TMAO, +85 -> +68 *m*/*z* (18 V); for choline, +104-> +60/45 *m*/*z* (14/16 V); for d_4_-choline, +108 -> +60 *m*/*z* (15 V); for γ-butyrobetaine, +146 -> +60/87 *m*/*z* (12/12 V); for carnitine, +162 -> +103/60 *m*/*z* (16/14 V); for d_3_-carnitine, +165 -> +103 *m*/*z* (16 V); for betaine, +118 -> +59/58 *m*/*z* (16/16 V); for d_9_-betaine, +127 -> +68 *m*/*z* (16 V).

### Kinome screen, *K_d_*

TMA was assessed using KdELECT screening service (DiscoveRx) as described previously^35,36^. This technique is based on a competition-binding assay that quantitatively measures the ability of a compound to compete with an immobilized, active-site-directed ligand. The assay consists of DNA-tagged kinase, immobilized ligand, and the potent inhibitor. The ability of TMA to compete with the immobilized ligand was measured by quantitative PCR of the DNA tag. The binding constant (*K_d_*) was then calculated from duplicate 11-point dose– response curve. Kinase interaction tree plots were generated using TREEspot™ Software Tool and reprinted with permission from KINOMEscan®, a division of DiscoveRx Corporation, © Discoverx corporation 2015.

### Kinase activity assays

The TMA IC_50_ on IRAK4 was determined using Kinexus kinase-inhibitor activity profiling service (Kinexus, Vancouver, Canada). Protein kinase assays (in duplicate) were performed at ambient temperature for 30 min in a final volume of 25 μL according to the following assay reaction recipe:

***Component 1***. 5 μl of diluted active IRAK4 target (recombinant, full length, expressed by baculovirus in Sf9 insect cells with an GST tag (SignalChem Catalogue 112 – 10G); ∼10-50 nM final concentration in the assay).

***Component 2***. 5 μl of stock solution of substrate (Myelin Basic Protein 1mg/ml diluted in H_2_O).

***Component 3***. 5 μl of kinase assay buffer (25 mM MOPS, pH 7.2, 12.5 mM β-glycerol-phosphate, 25 mM MgCl_2_, 5 mM EGTA, 2 mM EDTA, 0.25 mM DTT, added just prior to assay initiation).

***Component 4***. 5 μl of compound (various concentration as indicated) or 10% DMSO for blank.

***Component 5***. 5 μl of ^32^P-ATP (250 μM stock solution, 0.8 μCi, Perkin Elmer).

The assay was initiated by the addition of ^32^P-ATP and the reaction mixture incubated at ambient temperature for 30 min. After the incubation period, the assay was terminated by spotting 10 μL of the reaction mixture onto Multiscreen phosphocellulose P81 plate. The Multiscreen phosphocellulose P81 plate was washed 3 times for approximately 15 min each in a 1% phosphoric acid solution. The radioactivity on the P81 plate was counted in the presence of scintillation fluid in a Trilux scintillation counter. Blank control was set up that included all the assay components except the addition of Myelin Basic Protein (replaced with equal volume of assay dilution buffer). The corrected activity for IRAK4 target was determined by removing the blank control value.

### Reagents

Glutamine (Glutamax, 35050061, Life Technologies), Fetal bovine serum (Life Technologies), crystal violet (C6158) and Trimethylamine (W324108) were from Sigma-Aldrich. Mouse IL-6 Quantikine ELISA kits (M6000B, R&D system), RNeasy Micro Kit (Qiagen). SuperScript II Reverse Transcriptase, IL6 Taqman probe Hs00174131_m1 and FAST master mix (Invitrogen). High Fat Diet (Special Diets Services) and Low Fat Diet (B & K Universal) were specifically formulated in^12^. Control Diet (D12450K; Research diet), LC-HFD (60% kcal fat and 20% kcal carbohydrates, D12492, Research diet), HC-HFD (60% kcal fat and 20% kcal carbohydrates with 17 g of choline/kg, D16100401i, Research diet). Mouse Insulin ELISA (10-1249-01, Mercodia). Isoflurane (10014451, Forene, Abbott). TriPure reagent (1667165, Roche). Reverse Transcription System kit (A3500, Promega). Mesa Fast qPCR (CS-CKIT-PROD, Eurogentec). MSD V-PLEX Plus Proinflammatory Panel 1 kit (K15048G Meso Scale Diagnostics).

### Statistics

Potential outliers were identified by a Grubbs test. For statistical comparisons between study groups, normality was tested using D’agostino-Pearson omnibus normality test, then one-way ANOVA was used, followed by Tukey’s *post hoc* testing when data were normally distributed, otherwise groups were compared using the two-tailed Mann-Whitney test (*P* < 0.05 considered to be statistically significant). Data are displayed as mean ± s.e.m in all figures. All cell culture experiments included at least three biological replicates (as indicated in figure legends). All animal cohorts included at least five animals in each study group (as indicated in figure legends) and animals were randomized to treatment groups.

## Extended Data Figure Legends

**Extended Data Figure 1.**
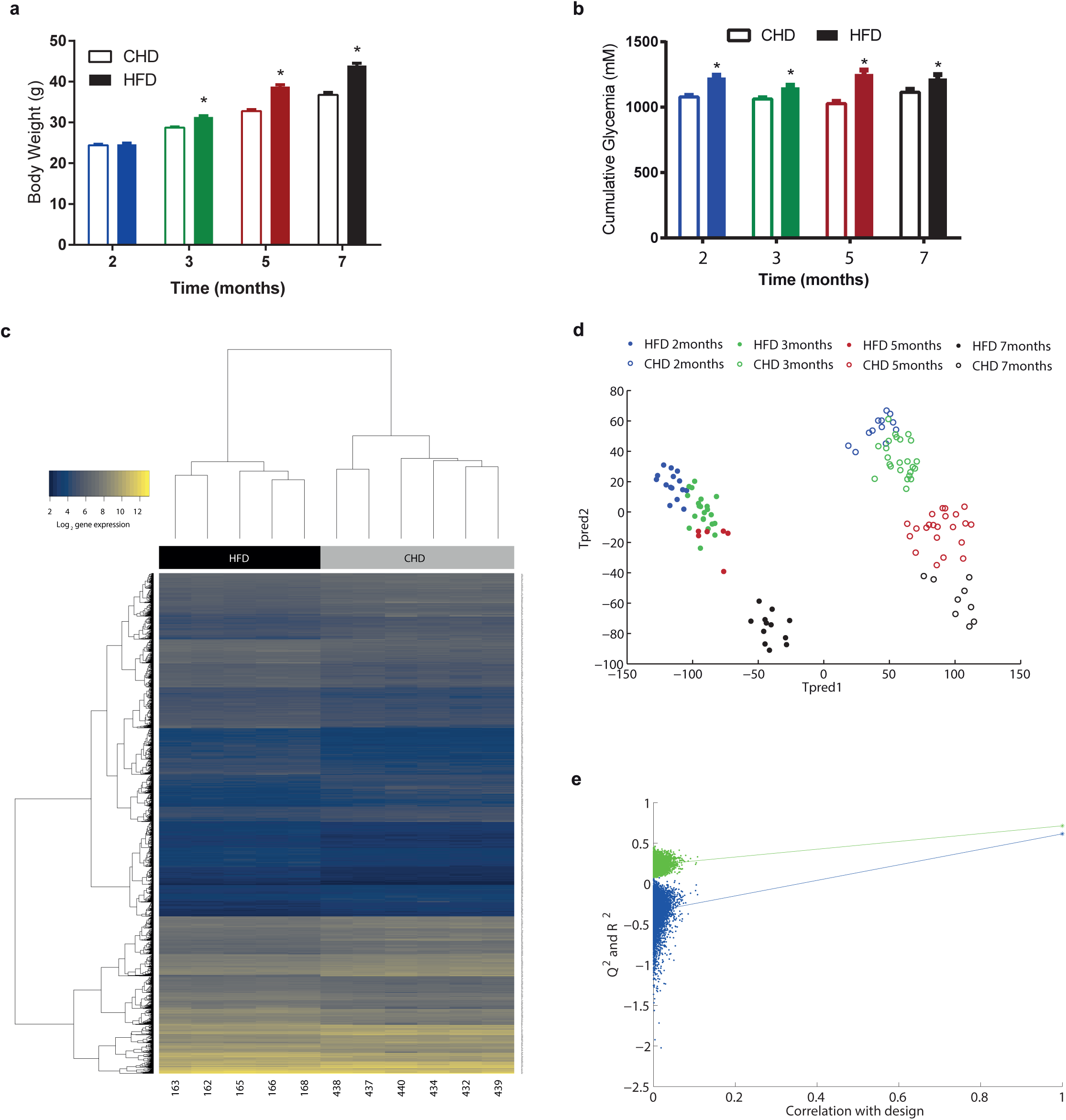
C57BL/6 mice fed HFD display low acute-phase gene expression in liver paralleled by high TMA excretion. Male mice were weaned at 3 weeks and fed CHD, before being randomly assigned to CHD group (n=270) or HFD (n=202) groups and monitored at 2 (72 CHD and 64 HFD), 3 (84 CHD and 57 HFD), 5 (68 CHD and 47 HFD) and 7 months (46 CHD and 34 HFD). (**a-b**) Effect of HFD-feeding on physiological parameters. (**a**) Body weight (BW) and (**b**) cumulative glycemia during GTT (AUC). (**c**) Heatmap of significantly (FDR < 0.1) differentially expressed liver genes after 4 months of HFD. (**d**) ^1^H-NMR-based metabotyping scores plot from a cross-validated O-PLS-DA model segregating the eight groups of mice according to both diet and age. (**e**) Empirical assessment of the significance of O-PLS goodness-of-fit parameters by generating a null distribution with 10,000 random permutations. Data are means ± s.e.m. Two-sided Mann-Whitney test (\**P* < 0.05 vs control). For the O-PLS-DA permutation test, the horizontal axis corresponds to the correlation between the original class membership variable (single dots on the right) and the randomly permuted class membership vectors (no longer correlated with the original class membership) (swarm of permutated models on the left). The *y* axis corresponds to goodness-of-fit parameter *R*^2^ (in green) derived for each O-PLS-DA model and the goodness-of-prediction parameter *Q*^2^ (in blue) derived by 7-fold cross-validation of each O-PLS-DA model. The *R*^2^ and *Q*^2^ parameters obtained from the original model in the top right corner are outside the confidence interval of the 10,000 randomly permuted models and therefore confirming the significance of the fitness and prediction capacity of the original O-PLS-DA model.

**Extended Data Figure 2.**
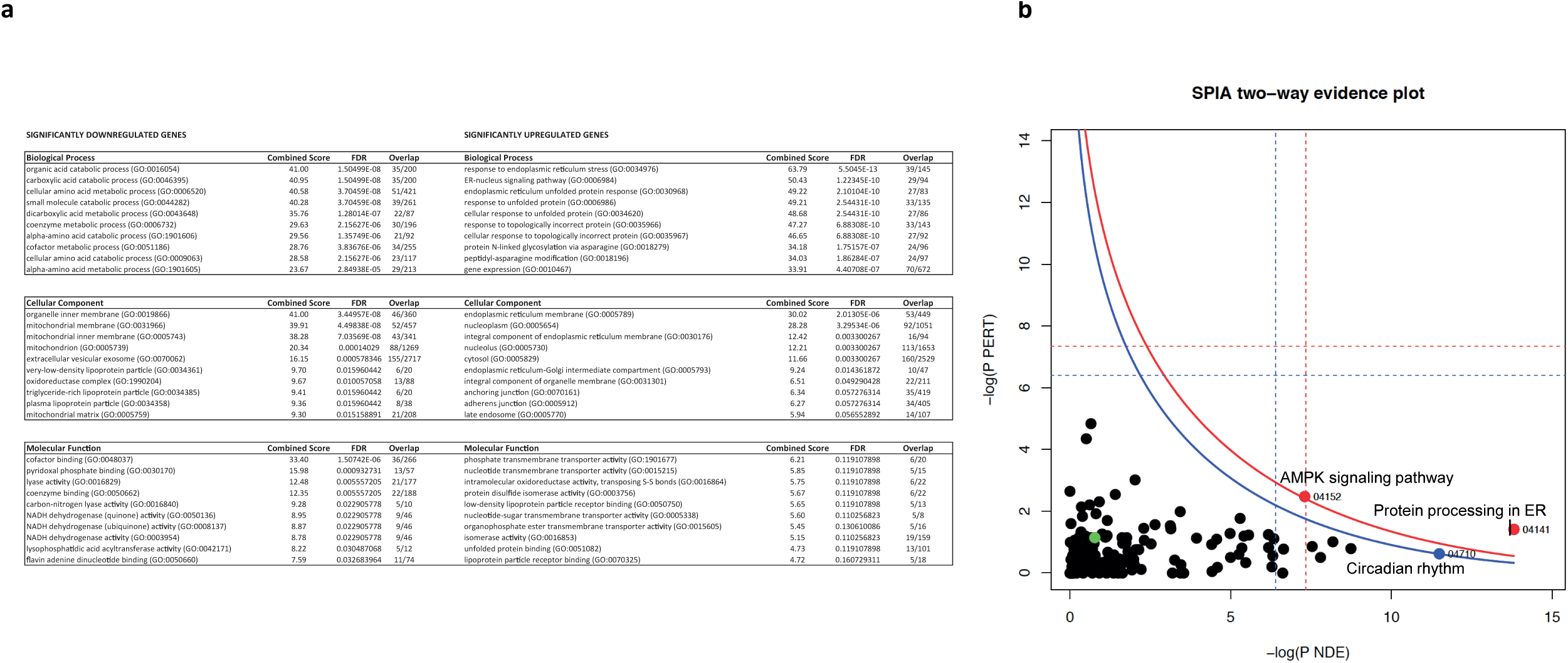
Pathway analysis for hepatic significant genes. (**a**) Gene ontology analysis for down- and up-regulated HFD-responsive genes. (**b**) Signaling pathway impact analysis for HFD-responsive genes. Pathways respectively on the right-hand side of the red and blue curves are significant after Bonferroni correction of the global p-values (pG) obtained by combining the pPERT and pNDE using the normal inversion method, and significant after a FDR correction of pG. Significant KEGG pathways: 04141, Protein processing in the endoplasmic reticulum; 04710, Circadian rhythm; 04152, AMPK signaling pathway.

**Extended Data Figure 3.**
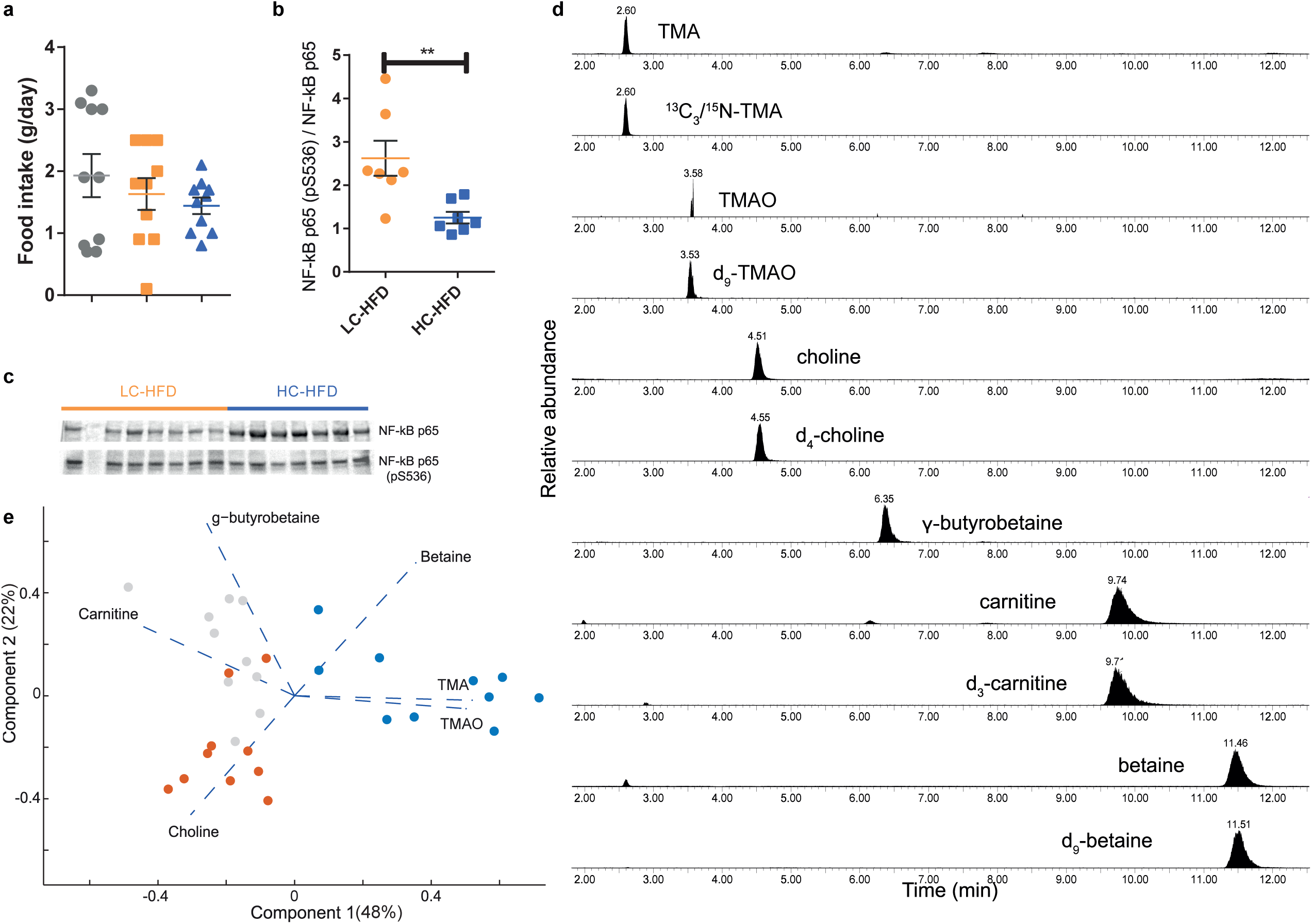
Choline supplementation corrects HFD adverse effects and associated methylamine measurements. (**a**) Food intake measured on the three tested diets (CHD, LC-HFD and HC-HFD). (**b-c**) Western blot analysis of liver NF-kB phosphorylation state between LC-HFD and HC-HFD. (**d**) Plasma sample chromatogram for the simultaneous detection of TMA, TMAO, choline, γ-butyrobetaine, carnitine, betaine and their isotopically labelled analogues. (**e**) Principal Component Analysis Biplot of plasma methylamine quantifications by UPLC/MS-MS from mice fed CHD, LC-HFD and HC-HFD, each component’s explained variance is shown in parenthesis. Data are means ± s.e.m. Two-sided Mann-Whitney test (\**P* < 0.05 vs control).

**Extended Data Figure 4.**
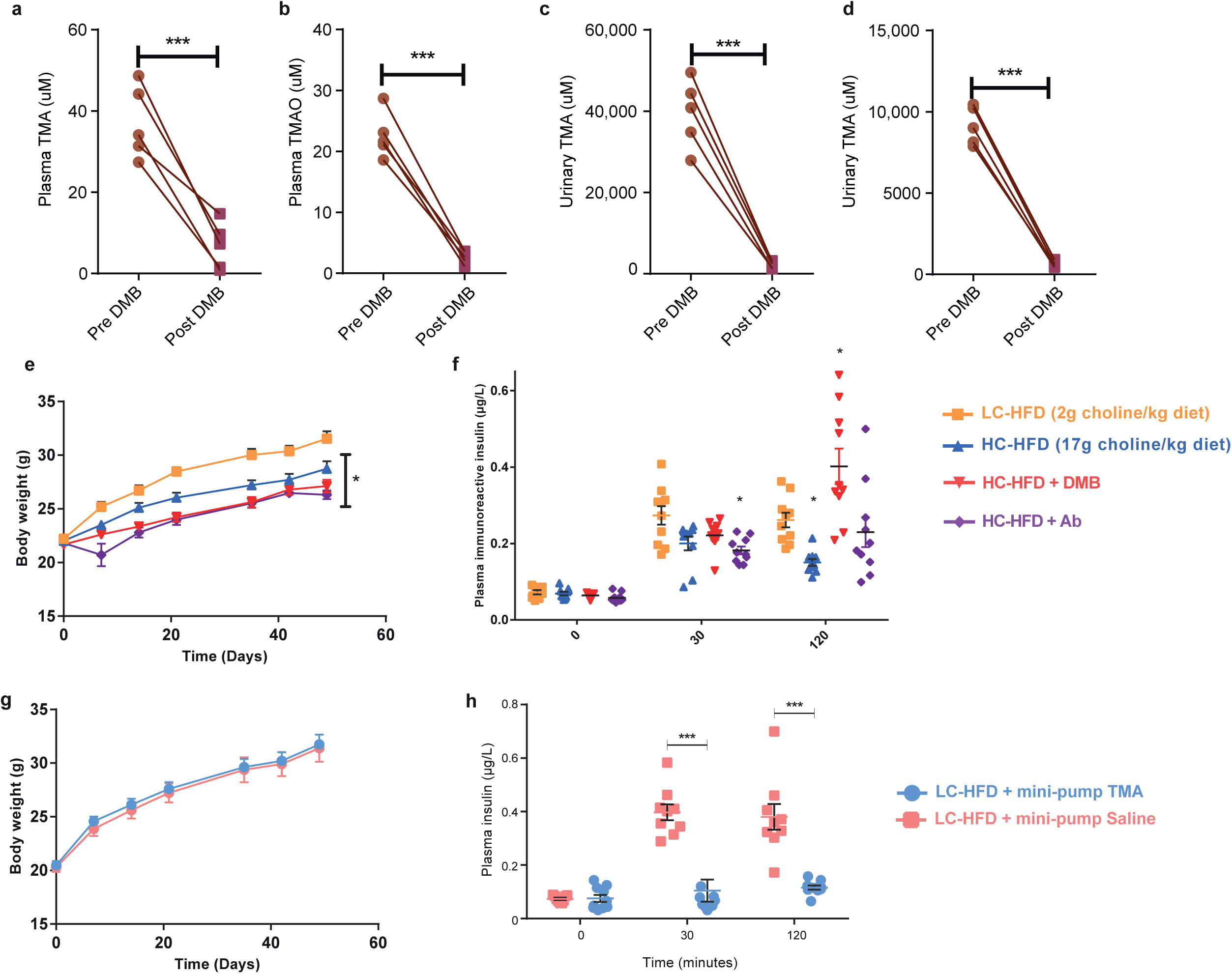
Modulation of TMA production and effects. (**a-d**) Confirmation of blockage of production of TMA (**a**, **c**) and TMAO (**b**, **d**) by 1% DMB measured in plasma (**a-b**) and urine (**c-d**). (**e**) Body weight monitoring during the whole experiment of mice fed a LC-HFD and a HC-HFD supplemented or not with DMB or treated with antibiotics. (**f**) Plasma insulin concentration during GTT at 0, 30 and 120 min post glucose injection. (**g**) Body weight monitoring during the whole experiment of mice fed a LC-HFD and chronically treated with TMA. (**h**) Plasma insulin concentration during GTT at 0, 30 and 120 min post glucose injection. Data are means ± s.e.m. Two-sided Mann-Whitney test (\**P* < 0.05 vs LC-HFD control).

**Extended Data Figure 5.**
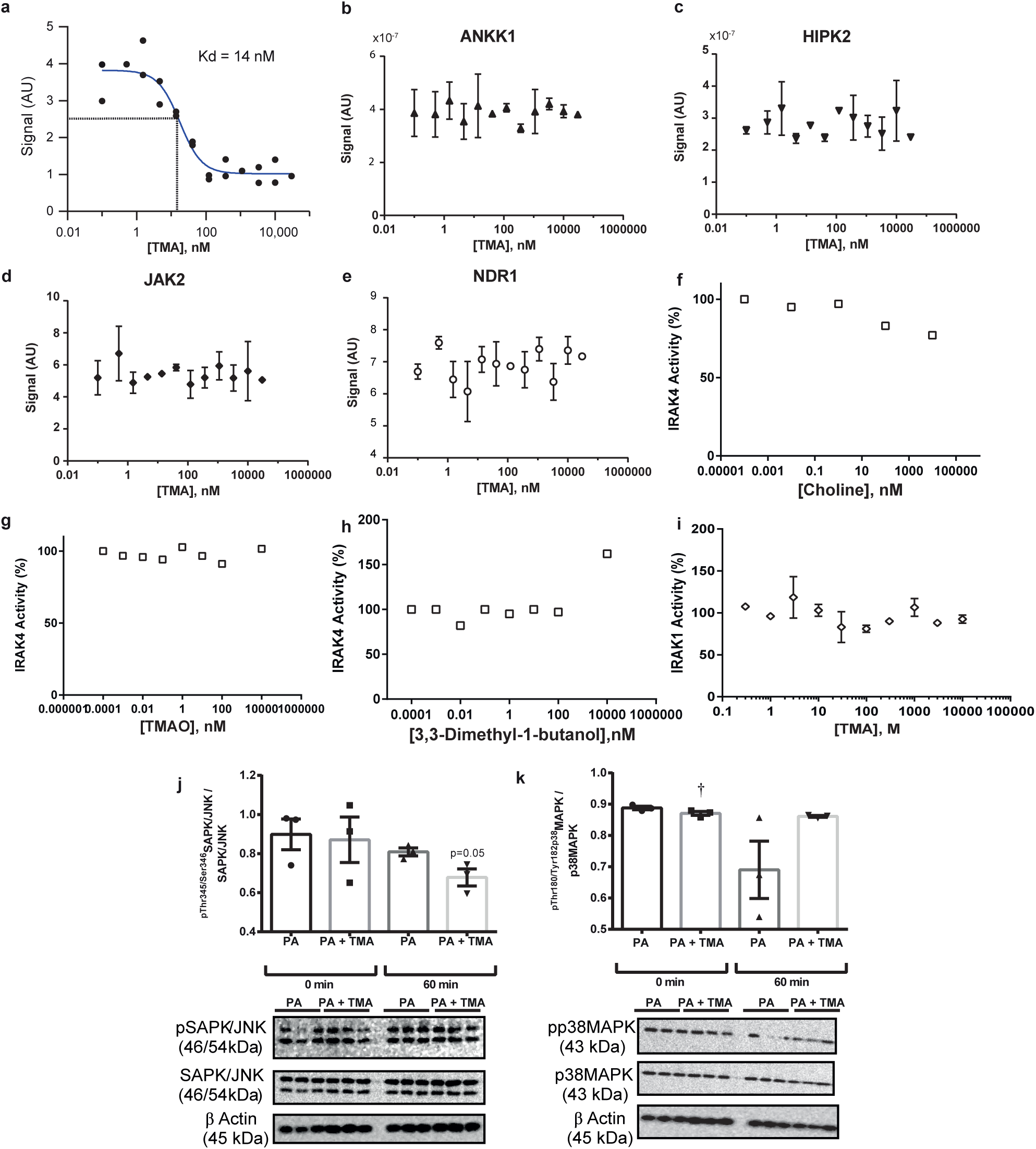
TMA does not bind with kinases ANKK1, HIPK2, JAK2, and NDR1 identified as candidates in the kinome screen, but makes significant contact with IRAK4. (**a**) Confirmation of a physical interaction between TMA and IRAK4 using TMA at concentrations ranging from 0.1 nM to 100 µM, resulting in a dissociation constant (*K_d_*) of 14 nM. (**b-e**) Physical interaction test between TMA and the four hits identified using a kinome screen, using TMA concentrations ranging from 0.1 nM to 100 µM for (**b**) ANKK1, (**c**) HIPK2, (**d**) JAK2 and (**e**) NDR1. IRAK4 kinase activity is not significantly inhibited by choline (**f**), TMAO (**g**) or DMB (**h**), while TMA does not have any inhibitory effect on IRAK-1 kinase activity (**i**). (**j-k**) Phosphorylation for ^pThr183/Tyr185^SAPK/JNK/SAPK/JNK (**j**) and ^pThr180/Tyr182^p38MAPK/p38MAPK (**k**) ratios. Data are means ± s.e.m. One-way ANOVA followed by Tukey’s *post hoc* tests (superscript letters for factor levels *P*<0.05) on log-transformed data.

**Extended Data Figure 6.**
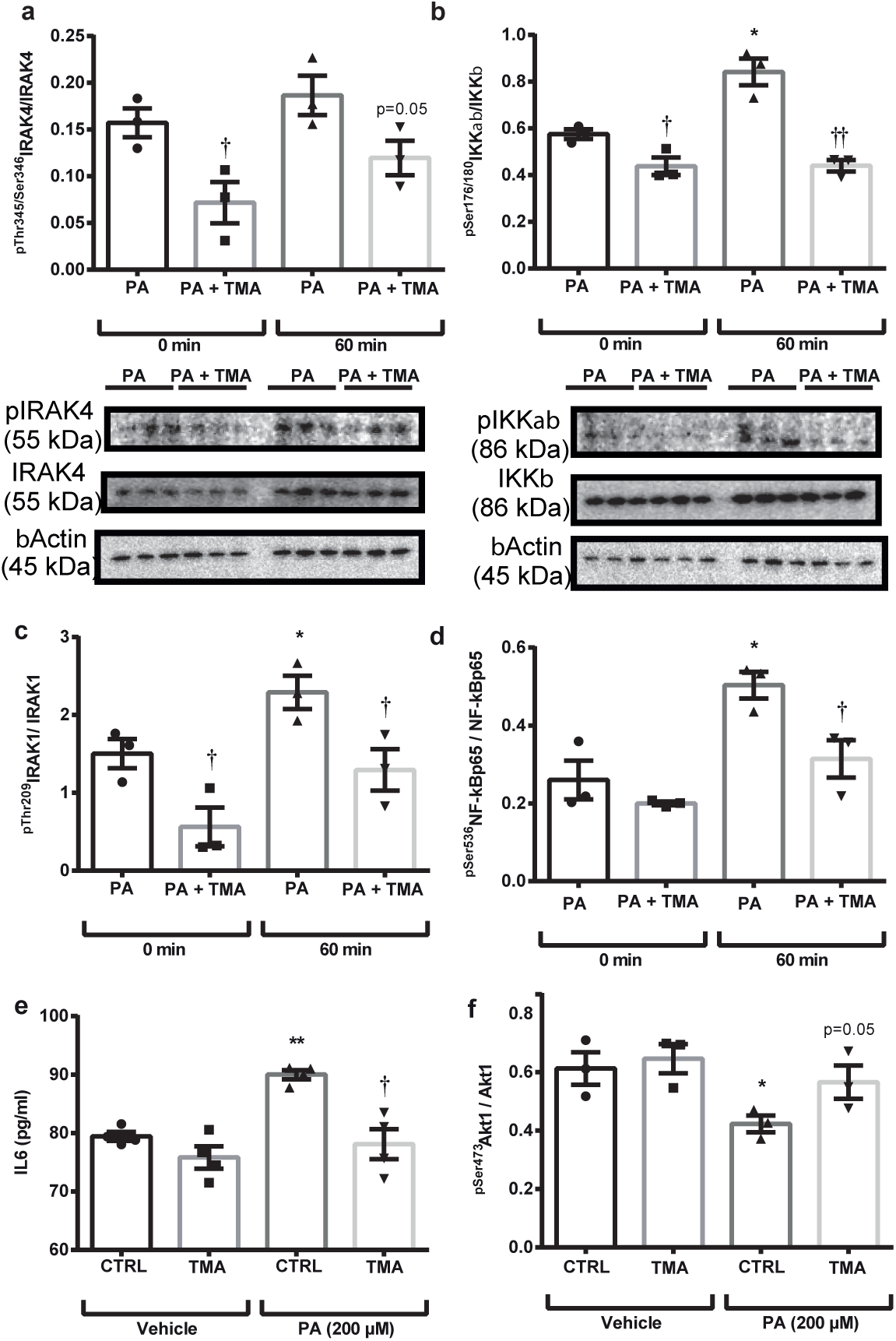
TMA inhibits IRAK4 and suppresses TLR4-mediated pro-inflammatory response in primary human hepatocytes. (**a-d**) Effect of palmitate (PA; 200 μM) administration and TMA pre-treatment (0.1 mM, 30 min) in primary human hepatocytes at 0 and 60 min on ^pThr345/Ser346^IRAK4/IRAK4 (**a**), ^pThr209^IRAK1/IRAK1 (**b**), ^pSer176/180^IKKαβ/IKKβ (**c**), ^pSer536^NF-κBp65/NF-κBp65 (**d**) ratios. *p<0.05 and **p<0.01 *vs* 0 min; ^†^p<0.05 and ^††^p<0.01 *vs* vehicle; ^b^p=0.07 *vs* vehicle. Effect of PA (200 μM) administration (4 h) with and without TMA pre-treatment (0.1 mM, 30 min) on IL6 accumulation in hepatocyte media (**e**) and on ^pSer473^Akt1/Akt1 after insulin stimuli (100 nM, 10min) (**f**). *p<0.05 and **p<0.01 *vs* control-vehicle; †p<0.05 *vs* control-PA.

**Extended Data Figure 7.**
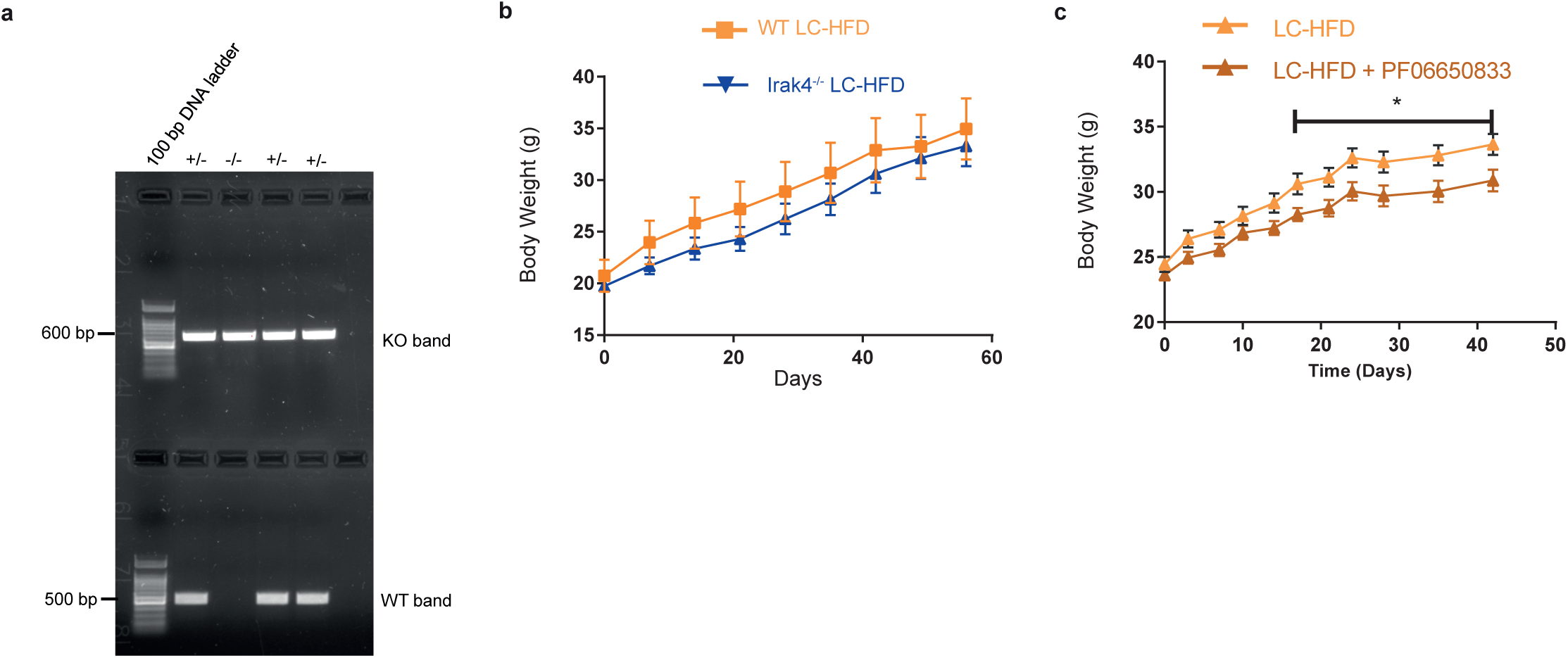
IRAK4 knock-out does not affect body weight while chronic treatment with PF06650833 reduces it in mice fed a low-choline HFD. (**a**) representative agarose gel of PCR products from +/-irak4 and -/-irak4 mice. (**b-c**) Body weight monitoring during the whole experiment of *Irak4*^-/-^ mice (**b**) or PF06650833-treated mice (**c**) fed a LC-HFD.

## Supplementary Tables

**Supplementary Table 1: Hepatic genes differentially expressed by HFD-feeding.** Hepatic transcriptomes were analysed using Limma^55^ and genes are considered significant if FDR<0.1.

**Supplementary Table 2: Down-regulated HFD-responsive hepatic pathways.** Gene ontology analysis was performed using Enrichr^56^ and pathways are considered significant if FDR<0.1.

**Supplementary Table 3: Up-regulated HFD-responsive hepatic genes.** Gene ontology analysis was performed using Enrichr^56^ and pathways are considered significant if FDR<0.1.

**Supplementary Table 4: Hepatic signalling pathways impacted by HFD.** Signalling pathway impact analysis was performed using SPIA^57^ and pathways are considered significant if pGFWER<0.1.

**Supplementary Table 5. TMA kinome screen.** Screening results for single-dose TMA binding to 456 kinases. “Percent control” corresponds to the percentage of DNA amplified by qPCR compared to the control condition. Values below 35 % (in green) are considered as positive hits.

**Supplementary Table 7. Diet formulations.** Specific fat, protein, carbohydrate and choline content in regimens formulated by Research Diet are summarized based on per cent w/w and per cent kcal.

